# AMPK-dependent phosphorylation is required for transcriptional activation of TFEB/TFE3

**DOI:** 10.1101/2021.01.27.428292

**Authors:** Mathieu Paquette, Leeanna El-Houjeiri, Linda C. Zirden, Pietri Puustinen, Paola Blanchette, Hyeonju Jeong, Kurt Dejgaard, Peter M. Siegel, Arnim Pause

**Author notes:** **Corresponding Author**, Arnim Pause.

## Abstract

Increased autophagy and lysosomal activity promote tumor growth, survival and chemo-resistance. During acute starvation, autophagy is rapidly engaged by AMPK activation and mTORC1 inhibition to maintain energy homeostasis and cell survival. TFEB and TFE3 are master transcriptional regulators of autophagy and lysosomal activity and their cytoplasm/nuclear shuttling is controlled by mTORC1-dependent multisite phosphorylation. However, it is not known whether and how the transcriptional activity of TFEB or TFE3 is regulated. We show that AMPK mediates phosphorylation of TFEB and TFE3 on three serine residues, leading to TFEB/TFE3 transcriptional activity upon nutrient starvation, FLCN depletion and pharmacological manipulation of mTORC1 or AMPK. AMPK loss does not affect TFEB/TFE3 nuclear localization nor protein levels but reduces their transcriptional activity. Collectively, we show that mTORC1 specifically controls TFEB/TFE3 cytosolic retention whereas AMPK is essential for TFEB/TFE3 transcriptional activity. This dual and opposing regulation of TFEB/TFE3 by mTORC1 and AMPK is reminiscent of the regulation of another critical regulator of autophagy, ULK1. Surprisingly, we show that chemoresistance is mediated by AMPK-dependent activation of TFEB, which is abolished by pharmacological inhibition of AMPK or mutation of serine 466/467/469 to alanine residues within TFEB. Altogether, we show that AMPK is a key regulator of TFEB/TFE3 transcriptional activity, and we validate AMPK as a promising target in cancer therapy to evade chemotherapeutic resistance.

## Introduction

Macroautophagy/autophagy, the process of cellular self-degradation, regulates many processes including energy supply during development and in response to acute nutrient stress [1,2]. Lysosomal biogenesis and autophagic activities are rapidly upregulated in response to starvation, oxidative stress, organelle damage and chemotherapy resistance [3]. Degradation of cargos inside the lysosome is a multistep process that is conserved in eukaryotes from yeast to multicellular organisms [4]. Fragments resulting from autophagic protein degradation are recycled to produce energy, allowing cells to survive under harsh circumstances. Deregulation of these pathways has been linked to various human disorders, including myopathy, neurodegeneration, cancer and metabolic syndrome [5,6].

The Microphthalmia family of bHLH-LZ (MiT/TFE) transcription factor family, namely Transcription Factor EB (TFEB) and Transcription Factor E3 (TFE3) [7] are master transcriptional regulators of autophagy and lysosomal biogenesis. TFEB and TFE3 homo- or hetero-dimerize and bind to gene promoter regions to activate a panel of genes involved in autophagy, lysosomal biogenesis, lipid metabolism, innate immune response and pathogen resistance [8–11]. They are redundant in some cellular contexts while indispensable in others [8,12,13].

TFEB and TFE3 have been linked with tumorigenesis. Both proteins have been implicated in chromosomal translocations resulting in the fusion of partial *TFEB/TFE3* coding regions with strong promoters, such as *MALAT1, PRCC, ASPSCR1, SFPQ, NONO* and *CLTC* [6,14]. Such fusion events enhance TFEB/TFE3 expression and activity and is strongly associated with juvenile renal cell carcinoma (RCC) and alveolar soft part sarcoma [14,15]. The oncogenic effect of these transcription factors is likely due to altered gene expression. Increased autophagy and lysosomal biogenesis may activate several pathways promoting cell survival, tumor growth and progression [2]. Typically, autophagy is induced during cancer therapy, which protects cancer cells and leads to drug resistance and refractory cancer [16]. Therefore, reducing TFEB/TFE3 activity might be important in cancer therapy settings.

Interestingly, TFEB and TFE3 also play important roles in non-dividing neurons to clear toxins and misfolded proteins through regulation of autophagic pathways [17]. Neurodegenerative diseases, including Alzheimer disease, Parkinson disease, and Huntington disease (HD) are characterized by the accumulation of intracellular aggregates in the brain [18]. Specifically in HD, expansion of a CAG trinucleotide repeat in the first exon of the Huntingtin (HTT) gene generates a protein containing an expanded polyglutamine (polyQ) tract, leading to pathogenic misfolding [17]. Increased autophagic/lysosomal activity was shown to reduce aggregates, revert symptoms and to restore cognitive capabilities [19–22]. Understanding the pathways modulating TFEB/TFE3 activity is therefore crucial to identify new targets for treatment of autophagy and lysosomal biogenesis related diseases, such as Alzheimer disease, Parkinson disease and HD.

To date, the best characterized regulator of TFEB/TFE3 is the serine/threonine kinase mechanistic Target of Rapamycin Complex 1 (mTORC1). mTORC1 is a mediator of cellular growth and proliferation that primarily integrates stress and growth signals by phosphorylating downstream targets to promote anabolic processes such as protein synthesis [23]. Under nutrient replete conditions, mTORC1 negatively regulates catabolic processes, including autophagy, by directly phosphorylating and inhibiting TFEB (serine residues 211 and 142) and TFE3 (serine residue 321). These phosphorylation events inhibit TFEB and TFE3 activation by promoting their cytoplasmic retention [24–27]. Conversely, under nutrient deficient conditions, these repressive phosphorylation events are removed, resulting in the nuclear translocation of TFEB/TFE3 and activation of autophagy and lysosomal biogenesis [8,11,28]. Similarly, mTORC1 also inhibits autophagy by direct phosphorylation of Unc-51-Like Kinase 1 (ULK1) under nutrient replete conditions [29–31].

AMP-activated protein kinase (AMPK) activates TFEB by blocking the activity of mTORC1 [32] and by increasing the levels of the Coactivator-Associated Arginine Methyltransferase (CARM1), an important cofactor for TFEB activity [33]. AMPK is an energy sensor and plays an essential role in the control of cellular bioenergetics [34] by responding to various stresses including those that induce changes in the cellular AMP:ATP ratio or modulation in intracellular calcium [35]. Upon its activation, AMPK maintains energy homeostasis by promoting energy production and limiting energy expenditure [35]. AMPK activation negatively regulates mTORC1 in two different ways. First, through direct phosphorylation and activation of the Tuberous Sclerosis Complex 1 and 2 (TSC1/2), AMPK inactivates the Ras homolog enriched in the brain (RHEB) protein, leading to subsequent mTORC1 inhibition [36]. Second, through phosphorylation of the Regulatory Associated Protein of mTOR (RAPTOR), AMPK promotes the binding of the chaperones 14-3-3 and mTORC1 inactivation [37]. Moreover, AMPK also regulates autophagosome maturation and autophagy by direct phosphorylation of ULK1 [29–31].

We and others have shown that AMPK can be chronically activated by the loss of Folliculin (FLCN) or loss of FLCN Interacting Proteins (FNIP1 and FNIP2) [38–44]. FLCN was also identified as a negative regulator of TFEB and TFE3 nuclear localization and activity [45,46]. *FLCN* is the gene associated with the Birt-Hogg-Dubé neoplastic syndrome in humans, characterized by predisposition to renal cell carcinoma (RCC), skin fibrofolliculoma, lung cysts and increased risk for spontaneous pneumothorax [47]. FLCN is thought to activate mTORC1 through its GTPase activating protein activity (GAP) toward the Ras-related GTPase C and Ras-related GTPase D (RagC/D) upon nutrient replenishment [48,49]. RagC/D, and the other members Ras-related GTPase A and Ras-related GTPase B (RagA/B), must be in an active conformation to anchor mTORC1 at the lysosome and to allow sensing of amino acids [50]. Therefore, FLCN loss leads to inhibition of RagC/D and removal of mTORC1 from the lysosomal surface, reducing phosphorylation and cytoplasmic sequestration of TFEB/TFE3 [51–53].

In this study, we assessed the role of AMPK in the regulation of TFEB/TFE3 activity under several conditions of cellular stress. We showed that although the sub-cellular localization of TFEB/TFE3 was regulated by mTORC1, AMPK was required for increased TFEB/TFE3 transcriptional activity, since deletion of AMPK completely blocked TFEB/TFE3 activity upon nutrient starvation, inhibition of mTORC1 or deletion of FLCN. We identified AMPK-dependent phosphorylation sites within TFEB in a highly conserved serine cluster (S466, S467 and S469), which were dispensable for proper sub-cellular TFEB localization but required for its transcriptional activity. Serine to alanine substitution mutations were sufficient to inhibit TFEB transcriptional activity in the nucleus, even in the context of mTORC1 inhibition or in presence of a constitutively nuclear TFEB mutant (S142A/S211A), which cannot be phosphorylated by mTORC1. AMPK-dependent phosphorylation of TFEB was required for increased resistance to the chemotherapy drug doxorubicin. Our findings shed new light on the interplay of two major metabolic kinases, AMPK and mTORC1, which both regulate autophagy and lysosomal biogenesis via TFEB/TFE3.

## Results

### AMPK is required for TFEB/TFE3 activity

We first studied the differential TFEB/TFE3 sub-cellular localization status in wild-type (WT) mouse embryonic fibroblasts (MEFs) compared to AMPKα1/α2 double knock-out (AMPK DKO) MEFs in a nutrient starvation context, using Earle’s Balanced Salt Solution (EBSS). As a positive control, we used the allosteric AMPK activator AICAR. EBSS starvation for 2 h induced nuclear translocation of TFEB and TFE3 in both WT and AMPK DKO MEFs, while 2 h AICAR treatment induced nuclear translocation only in WT cells (Fig. 1A, quantified in B). EBSS inhibited mTORC1 signaling, as shown by reduction in the phosphorylated forms of the known downstream target ribosomal protein S6 (S6) in both WT and AMPK DKO cells (Fig. 1C), suggesting that mTORC1 inhibition upon EBSS starvation was independent of AMPK. Indeed, mTORC1 can directly sense levels of amino acids independently of AMPK activity [50,54–56]. Nonetheless, AMPK was activated by EBSS starvation as shown by the increase in its phosphorylated form (p-AMPK) and phosphorylation of its known downstream target Acetyl-CoA carboxylase 1 or 2 (ACC1/2) (Fig. 1C). AICAR also activated AMPK and inhibited mTORC1 in WT cells, but not in AMPK DKO cells, pointing to a specific action of AMPK (Fig. 1D). We did not observe a reduction in TFEB/TFE3 protein levels upon above mentioned treatments or in AMPK DKO cells when assessed by fluorescence intensity or western blot. Surprisingly, EBSS-dependent increases in DQ-BSA fluorescence intensity, which measures lysosomal protease activity and serves as a proxy for autophagosome/lysosome activity [32,57], was significantly abrogated in AMPK DKO cells (Fig. 1E, quantified in F) despite evident TFEB/TFE3 nuclear localization upon EBSS starvation. DQ-BSA is a quenched fluorogenic substrate for intracellular lysosomal proteases. Cleavage of the fluorescent peptides within the lysosome relieves quenching, resulting in a fluorescent signal. As expected, AICAR-dependent increase in DQ-BSA fluorescence intensity was abrogated in AMPK DKO cells (Fig. 1E, quantified in F). Treatment with EBSS and AICAR also increased the number of lysosomes in WT but not AMPK DKO cells, as assessed by counting the average number of Lysosomal-associated membrane protein 1 (LAMP1) puncta per cell (Fig. 1G). This increase in number was not accompanied by a difference in lysosome size (Fig. 1H). Moreover, we measured TFEB/TFE3 transcriptional activity using a luciferase reporter containing the TFEB/TFE3 consensus promoter region (Coordinated Lysosomal Expression and Regulation; CLEAR). Treatment of WT MEFs with EBSS and AICAR induced expression from the CLEAR luciferase reporter, which was not observed in AMPK DKO cells (Fig. 11, J). Furthermore, the induction of known TFEB/TFE3 target genes such as *Atp6v1c1, Atp6v0d1, Ctsa, Ctsd, Gabarap* and *Sdha* by EBSS starvation and AICAR treatment was strongly reduced in AMPK DKO cells as measured by Reverse transcription quantitative polymerase chain reaction (RT-qPCR) (Fig. 1K, L). Taken together, these results suggest that although TFEB/TFE3 nuclear localization is not dependent on AMPK upon EBSS starvation, AMPK plays an essential role in regulating TFEB/TFE3 transcriptional activity.

**Figure 1.**
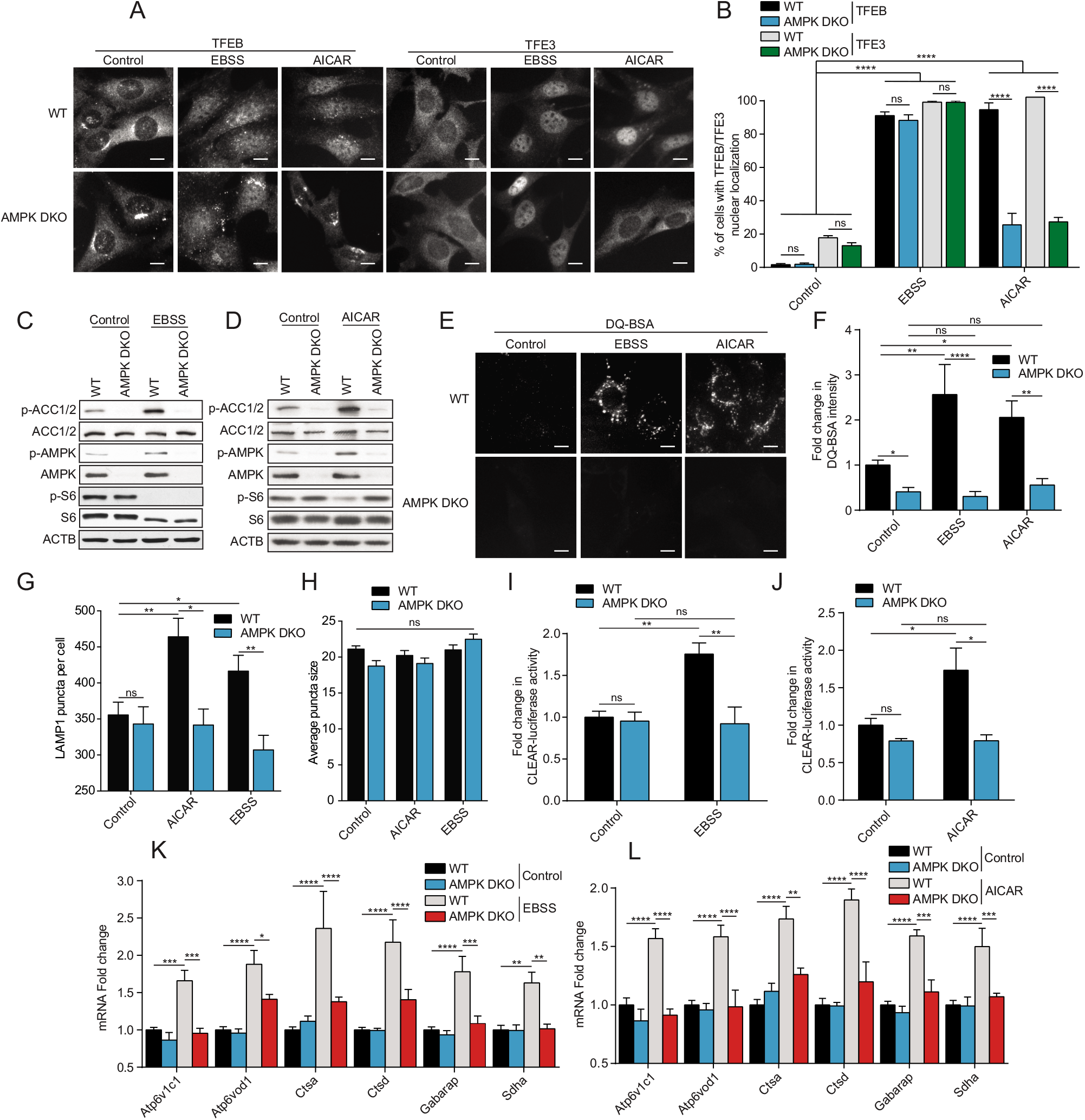
AMPK is required for complete TFEB/TFE3 activity upon EBSS starvation and AICAR treatment. (**A**) WT and AMPK DKO MEFs were incubated in complete media (Control), starved (EBSS) or in presence of AICAR (2 mM) for 2 h. Cells were fixed, permeabilized, and stained with an antibody against TFEB or TFE3. Scale bar, 20 μm. Images are representative of three independent experiments. (**B**) Quantification of the percentage of indicated MEF lines with nuclear TFEB or TFE3 upon treatments as indicated in (A) (mean ± SEM of three independent experiments, two-way ANOVA, ns=not significant, ****P < 0.0001; n > 200 cells per condition). (**C-D**) Immunoblot of protein lysates of indicated MEF lines treated as indicated in (A). Data are representative of three independent experiments. (**E**) Representative images of indicated MEF lines after 1 h of incubation with DQ-BSA-Red followed by a 2 h chase in complete media (Control), starved (EBSS) or in presence of AICAR (2 mM) prior to fixation. Scale bar, 20 μm. Images are representative of four independent experiments. (**F**) Relative lysosomal activity, as determined by DQ-BSA assay, in indicated MEF lines upon treatment as indicated in (E) (mean ± SEM of four independent experiments, two-way ANOVA, ns=not significant, *P< 0.05, **P < 0.01, ****P < 0.0001; n > 200 cells per condition). (**G**) Quantification of the number of LAMP1 puncta upon treatment as indicated in (A). (mean ± SEM of three independent experiments, two-way ANOVA, ns=not significant, *P < 0.05, **P < 0.01; n > 50 cells per condition). (**H**) Quantification of the average LAMP1 puncta size upon treatment as indicated in (A). (mean ± SEM of three independent experiments, two-way ANOVA, ns=not significant, n > 50 cells per condition). (**I-J**) Relative TFEB/TFE3 transcriptional activity, as determined by CLEAR-luciferase promoter activity normalized against CMV-Renilla in indicated MEF lines upon treatment as indicated in (A) (mean ± SEM of the luminescence fold change from five independent experiments, two-way ANOVA, ns=not significant, *P < 0.05, **P < 0.01). (**K-L**) Relative quantitative real-time PCR analysis of TFEB/TFE3 target genes mRNA transcript levels in indicated MEF lines upon treatment as indicated in (A) (mean ± SEM of the RNA fold change of indicated mRNAs from five independent experiments, two-way ANOVA, *P < 0.05, ***P< 0.001, ****P < 0.0001).

### mTORC1 inhibition requires AMPK for complete TFEB/TFE3 activity

We showed above that EBSS starvation of WT cells led to simultaneous AMPK activation and mTORC1 inhibition. We also observed mTORC1 inhibition upon EBSS starvation in AMPK DKO cells, which is probably due to the ability of mTORC1 to sense amino acid levels independently of AMPK. Since AMPK is a negative upstream regulator of mTORC1, it is conceivable that the observed inhibition of mTORC1, and not the activation of AMPK, led to the translocation of TFEB/TFE3 in both cases. To test this hypothesis, we examined whether mTORC1 inhibition using Torin1 would show a similar AMPK-dependent effect on TFEB/TFE3 transcriptional activity. To extend our observations beyond MEF cells, we generated human HEK293T AMPK DKO using CRISPR-Cas9. Gene knockout was confirmed by western blot using an antibody recognizing both the α1 and α2 subunits (Fig. 2A). Torin1 treatment for 2 h led to inhibition of mTORC1 signaling in both MEFs and HEK293T WT and AMPK DKO cell lines (Fig. 2A, B), as shown by a decrease in levels of pS6, and led to nuclear localization of TFEB/TFE3 in both WT and AMPK DKO cells (Fig. 2C, quantified in D and E). Since TFEB is not well expressed in HEK293T cells we focused only on TFE3 localization in these cells. In line with results in Figure 1, whereas Torin1 treatment led to an increase in lysosomal activity measured by DQ-BSA fluorescence in WT cells, this effect was abolished in AMPK DKO cells (Fig. 2F, quantified in G). This was accompanied with an increase in lysosome number in WT cells compared to AMPK DKO when treated with Torin1 (Fig. 2H), but no statistical difference in terms of lysosome size was observed (Fig. 2I). In addition, TFEB/TFE3 transcriptional activity was induced upon treatment with Torin1, which was not observed in AMPK DKO cells by both CLEAR luciferase reporter (Fig. 2J, K) and RT-qPCR (Fig. 2L, M) assays in MEFS and HEK293T cells. These results suggest that TFEB/TFE3 transcriptional activity is dictated by an mTORC1-independent but AMPK-dependent mechanism, whereas nuclear translocation is only dependent on mTORC1.

**Figure 2.**
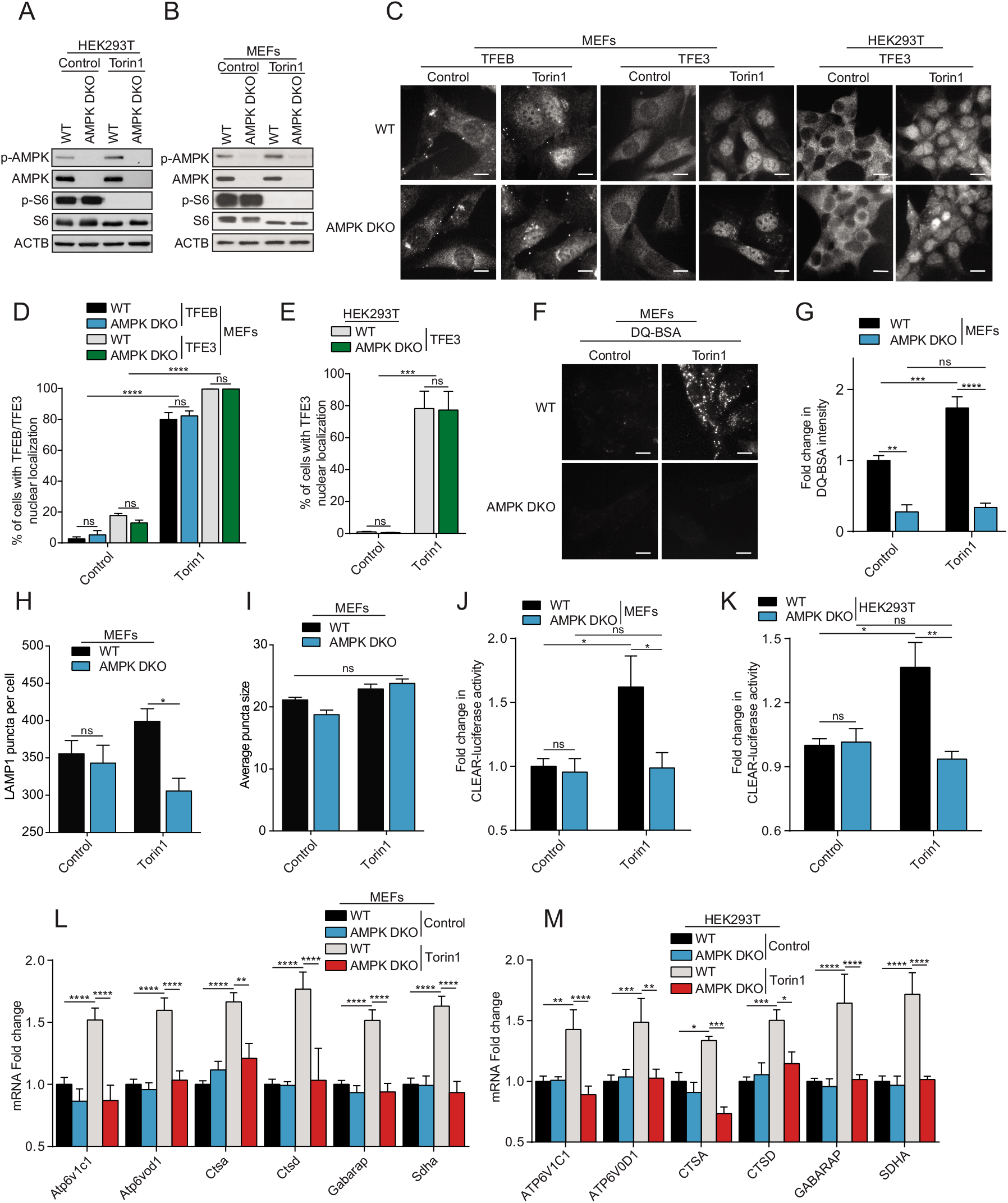
mTORC1 inhibition requires AMPK for complete TFEB/TFE3 activity. (**A-B**) Immunoblot of protein lysates of HEK293T (A) and MEFs (B) with indicated genotypes incubated in presence of DMSO (Control) or Torin1 (1 μM) for 2 h. Data are representative of three independent experiments. (**C**) Representative images of cell lines treated as indicated in (A-B). Cells were fixed, permeabilized, and stained with an antibody against TFEB or TFE3. Scale bar, 20 μm. Images are representative of three independent experiments. (**D-E**) Quantification of the percentage of MEFs (D) and HEK293T (E) with nuclear TFEB or TFE3 upon treatments as indicated in (A-B) (mean ± SEM of three independent experiments, two-way ANOVA, ns=not significant, ***P < 0.001, ****P < 0.0001; n > 200 cells per condition). (**F**) Representative images of MEFs after 1 h of incubation with DQ-BSA-Red followed by a 2 h chase in complete media in presence of DMSO (Control) or Torin1 (1 μM) prior to fixation. Scale bar, 20 μm. Images are representative of four independent experiments. (**G**) Relative lysosomal activity, as determined by DQ-BSA assay, in MEFs upon treatment as indicated in (F) (mean ± SEM of four independent experiments, two-way ANOVA, ns=not significant, **P < 0.01, ***P < 0.001, ****P < 0.0001; n > 200 cells per condition). (**H**) Quantification of the number of LAMP1 puncta in MEFs upon treatment as indicated in (A). (mean ± SEM of three independent experiments, two-way ANOVA, ns=not significant, *P < 0.05; n > 50 cells per condition). (**I**) Quantification of the average LAMP1 puncta size in MEFs upon treatment as indicated in (A). (mean ± SEM of three independent experiments, two-way ANOVA, ns=not significant, n > 50 cells per condition). (**J-K**) Relative TFEB/TFE3 transcriptional activity, as determined by CLEAR-luciferase promoter activity normalized against CMV-Renilla in MEFs (J) and HEK293T (K) upon treatment as indicated in (A) (mean ± SEM of the luminescence fold change from four independent experiments, two-way ANOVA, ns=not significant, *P < 0.05, **P < 0.01). (**L-M**) Relative quantitative real-time PCR analysis of TFEB/TFE3 target genes mRNA transcript levels in MEFs (L) and HEK293T (M) upon treatment as indicated in (A) (mean ± SEM of the RNA fold change of indicated mRNAs from four independent experiments, two-way ANOVA, *P < 0.05, **P< 0.01, ***P < 0.001, ****P < 0.0001).

### AMPK is required for FLCN-dependent activation of TFEB/TFE3

To further substantiate our observations that AMPK is required for TFEB/TFE3 activity, we tested conditions where TFEB and TFE3 are constitutively active and localized to the nucleus, such as in FLCN KO cells [45,58]. To this end, we generated FLCN KO/AMPK DKO MEFs (TKO; Fig. 3A). Upon loss of FLCN, TFEB/TFE3 were localized permanently to the nucleus, which remained unchanged upon deletion of AMPK in TKO cells (Fig. 3B, quantified in C). These results demonstrate that the nuclear localization of these transcription factors, in the context of FLCN deletion, is independent of AMPK as we observed above with EBSS and Torin1 treatment. However, the transcriptional activity of TFEB/TFE3 observed in FLCN KO cells was abolished upon deletion of AMPK in TKO cells, as measured by DQ-BSA assay (Fig. 3D, quantified in E), CLEAR luciferase reporter assay (Fig. 3F) and RT-qPCR of known TFEB/TFE3 target genes (Fig. 3G), suggesting that AMPK is required for FLCN-dependent transcriptional activation of TFEB/TFE3.

**Figure 3.**
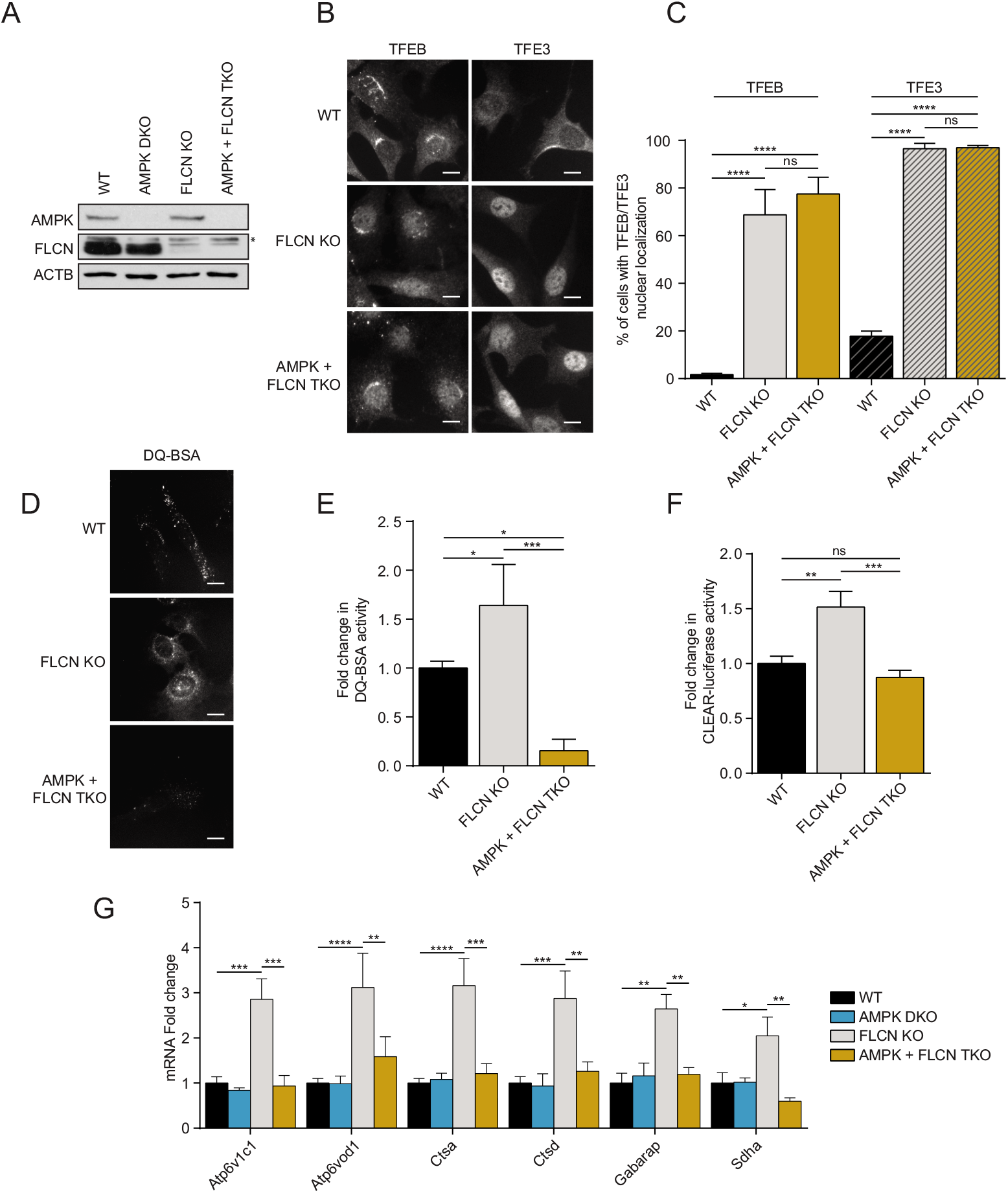
AMPK is required for FLCN-dependent activation of TFEB/TFE3. (**A**) Immunoblot of protein lysates of MEFs with indicated genotypes were incubated in complete media. Data are representative of three independent experiments. * represents non-specific bands. (**B**) Representative images of indicated MEF lines incubated in complete media. Cells were fixed, permeabilized, and stained with an antibody against TFEB or TFE3. Scale bar, 20 μm. Images are representative of three independent experiments. (**C**) Quantification of the percentage of nuclear TFEB or TFE3 in MEFs incubated in complete media (mean ± SEM of three independent experiments, one-way ANOVA, ns=not significant, ****P < 0.0001; n > 200 cells per condition). (**D**) Representative images of indicated MEF lines after 1 h of incubation with DQ-BSA-Red followed by a 2 h chase in complete media prior to fixation. Scale bar, 20 μm. Images are representative of three independent experiments. (**E**) Relative lysosomal activity, as determined by DQ-BSA assay in MEFs upon treatment as indicated in (D) (mean ± SEM of three independent experiments, one-way ANOVA, *P < 0.05, ***P < 0.001; n > 200 cells per condition). (**F**) Relative TFEB/TFE3 transcriptional activity, as determined by CLEAR-luciferase promoter activity normalized against CMV-Renilla in MEFs incubated in complete media (mean ± SEM of the luminescence fold change from six independent experiments, one-way ANOVA, ns=not significant, **P < 0.01, ***P < 0.001) (**G**) Relative quantitative real-time PCR analysis of TFEB/TFE3 target genes mRNA transcript levels in MEFs incubated in complete media (mean ± SEM of the RNA fold change of indicated mRNAs from four independent experiments, two-way ANOVA, ns=not significant, *P < 0.05, **P< 0.01, ***P < 0.001, ****P < 0.0001).

### AMPK is required for clearance of polyQ protein aggregates in a model of Huntington disease

To substantiate the physiological relevance of our results we used a cellular model of Huntington disease (HD), which is dependent on stimulation of autophagy and lysosomal biogenesis. Several groups have reported that mTORC1 inhibition leads to clearance of intracellular protein aggregates [59–61]. To this end, we generated stable WT, AMPK DKO, FLCN KO and AMPK/FLCN TKO MEFs expressing a doxycycline-inducible fusion protein consisting of the Cyan Fluorescent Protein (CFP) merged to a chain of 94 glutamine residues. Following a three-day induction with doxycycline, the polyQ chains form fluorescent aggregates mimicking aggregates formed in HD neurons that require autophagy and lysosomal activity for their clearance [62]. Treatment with AICAR or Torin1 led to a decrease in polyQ aggregates in WT cells but not in AMPK DKO cells (Fig. S1A, quantified in B). In addition, AMPK DKO cells displayed increased accumulation of polyQ aggregates upon treatment with vehicle consistent with impaired autophagic and lysosomal activity (Fig. S1A, quantified in B). Consistently with the constitutive TFEB/TFE3 nuclear localization and increased activity upon FLCN loss, FLCN KO cells displayed a reduction in protein aggregates (Fig. S1C, quantified in D). Interestingly, the additional AMPK loss in FLCN KO increased the number of cells with protein aggregates to a level similar to WT cells (Fig. S1C, quantified in D). Moreover, we observed a reduction of insoluble protein aggregates in WT cells but not in AMPK DKO as measured by fractionation of soluble versus insoluble protein (Fig. S1E, quantified in F). These results suggest that the presence and activity of AMPK is required for clearance of insoluble protein aggregates upon treatment with a mTORC1 inhibitor.

### AMPK phosphorylates TFEB and TFE3 on highly conserved serine cluster 466, 467, 469

Since AMPK is required for TFEB/TFE3 transcriptional activity and is a serine/threonine kinase, we next assessed whether AMPK regulates TFEB/TFE3 activity via phosphorylation. Since TFEB and TFE3 appear to be regulated in a similar fashion, we focused our phosphorylation studies on TFEB for simplicity. We developed an *in vitro* phosphorylation assay using recombinant AMPK and TFEB. After incubation with [γ-^32^P]ATP, we observed AMPK-dependent TFEB phosphorylation, which was enhanced after the addition of AMP, an allosteric AMPK activator (Fig. 4A). The level of phosphorylation on TFEB was similar to another well-characterized AMPK substrate, ULK1 (Fig. 4B). It is noteworthy to mention that some phosphorylation on ULK1 could be due to autophosphorylation. Strikingly, the phosphorylation efficiency of AMPK toward TFEB was more significant than towards ACC, suggesting that TFEB is a better substrate than this very well characterized substrate (Fig. 4C). Interestingly, immunoprecipitation of endogenous AMPK revealed that AMPK and TFEB form a complex in HEK293T cells, which was slightly increased upon the addition of AICAR (Fig. 4D). In order to reveal potential AMPK phosphorylation sites on TFEB, we purified recombinant fragments of GST-TFEB from bacteria, performed a kinase assay with purified AMPK, and analyzed GST-TFEB with mass spectrometry. This approach identified phosphorylation of the serine residues 467 and 469 (Fig. S2A-B). The C-terminal region is highly conserved throughout evolution and is maintained in other MiT/TFE family members (Fig. 4E). Moreover, these two serine residues and the adjacent serine (S466) resemble an AMPK phosphorylation motif (Fig. 4F). Although the phosphorylation motif is not a perfect fit, it has been previously shown that AMPK can potently phosphorylate its substrate without the canonical motif, such as ULK1 (Fig. 4F) [63–65]. Importantly, while GST-TFEB WT is efficiently phosphorylated by AMPK *in vitro*, a truncation of the 36 C-terminal residues led to a dramatic reduction in phosphorylation (Fig. 4G). Furthermore, although single point mutations of the three serine residues to alanine (S466, S467 or S469) did not reduce phosphorylation to a large extent, a triple S466/467/469A mutation led to a significant inhibition of AMPK-dependent TFEB phosphorylation *in vitro* (Fig. 4H). Similarly, point mutations of serine residues at 567, 568, 570 in TFE3 (corresponding to S466, S467, S469 in TFEB) to alanine also led to a significant reduction of AMPK-dependent TFE3 phosphorylation *in vitro* (Fig. 4I). Importantly, we generated a phosphospecific antibody raised against a triple-phosphorylated peptide p-S466/467/469 corresponding to the human C-terminal region of TFEB and p-S567/568/570 of TFE3 to measure potential phosphorylation by AMPK in cultured cells. This antibody recognized a phosphorylated TFEB fragment incubated in presence of purified AMPK, but not a fragment harboring serine to alanine mutations (Fig. 4J). As the phosphorylated peptide used to immunize the rabbits is perfectly conserved in TFE3, the antibody recognized a phosphorylated TFE3 fragment incubated in presence of AMPK as well, but not the fragment S567/568/570A (Fig. 4K). To assess potential phosphorylation by AMPK in cultured cells and to better discriminate between TFEB and TFE3, we immunoprecipitated both p-TFEB and p-TFE3 from HEK293T cells using this antibody and measured the relative abundance of each phosphoprotein using specific antibodies against total TFEB and TFE3. Starvation in EBSS revealed a 2.9-fold increase in the abundance of phosphorylated TFEB and a 2.6-fold increase in phosphorylated TFE3, which was not apparent in AMPK DKO cells (Fig. 4L, quantified in M). Similarly, treatment with AICAR increased the abundance of phosphorylated TFEB by 2.5-fold and TFE3 by 2-fold (Fig. 4L, quantified in M). Therefore, we conclude that AMPK phosphorylates TFEB and TFE3 on this highly conserved C-terminal serine cluster in vitro and in cultured cells.

**Figure 4.**
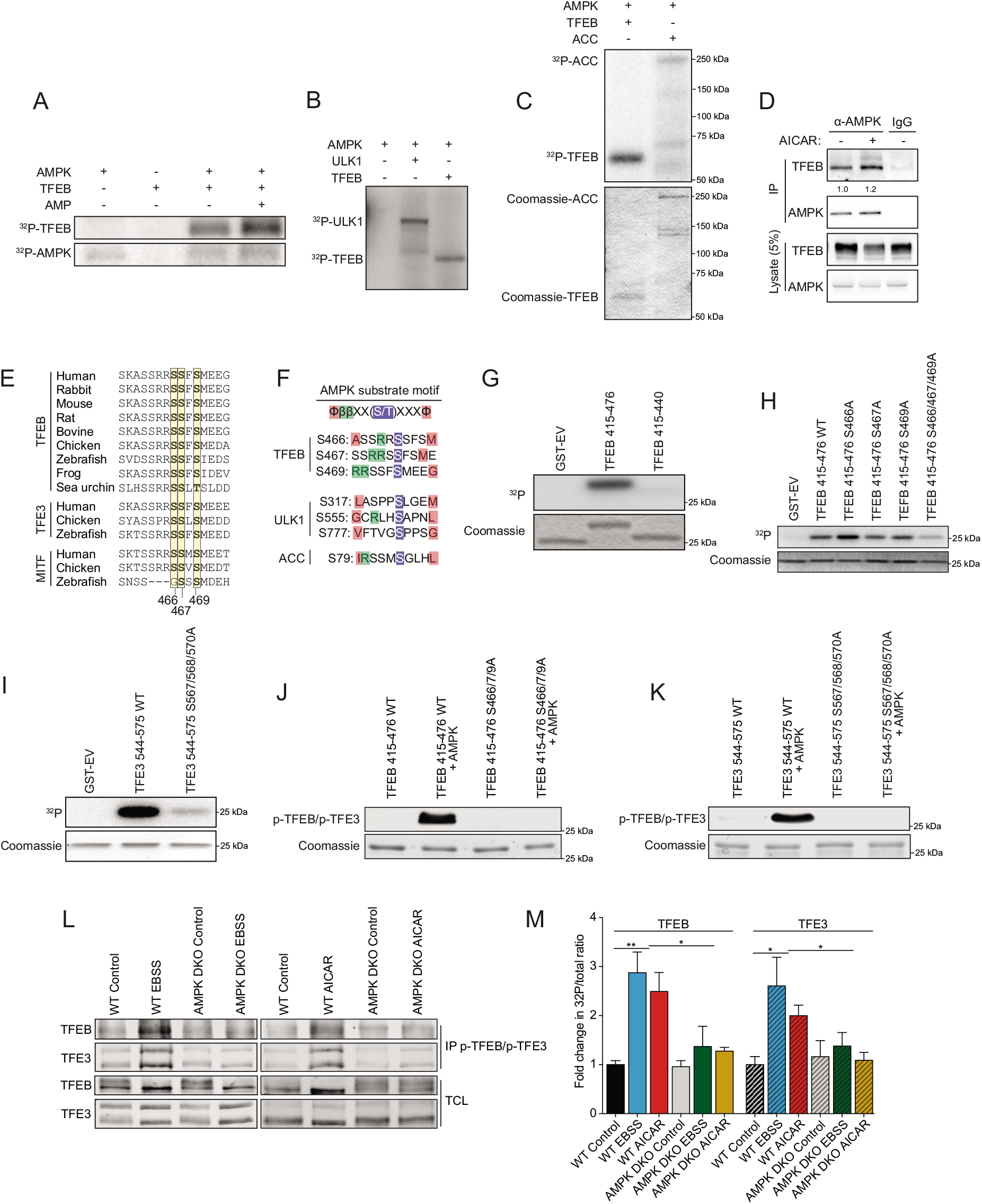
AMPK phosphorylates TFEB on S466, S467 and S469 *in vitro*. (**A**) *In vitro* kinase assays. Purified AMPK (0.1 μg) and recombinant TFEB (0.5 μg) were incubated with [γ-^32^P]ATP with or without AMP and analyzed by SDS-PAGE followed by autoradiography. Data are representative of three independent experiments. (**B**) *In vitro* kinase assays. Purified AMPK (0.1 μg), recombinant TFEB (0.5 μg), and recombinant ULK1 (0.5 μg) incubated as described in (A) (**C**) *In vitro* kinase assays. Purified AMPK (0.05 μg), recombinant TFEB (0.5 μg), and recombinant ACC (0.5 μg) incubated as described in (A). (**D**) Endogenous protein complexes were immuno-purified from HEK293T cells treated with vehicle or 2 mM AICAR for 2 h with AMPKα1/2 antibody and analyzed by immunoblotting with indicated antibodies. Non-specific IgG was used as a negative control. Data are representative of three independent experiments, quantification of TFEB protein levels normalized to AMPK protein level, Student’s t-test, p=0.03. (**E**) Alignment of TFEB/TFE3/MITF amino acid C-terminal regions from various species. (**F**) AMPK substrate motif alignment surrounding the serine residues 466, 467 and 469 in human TFEB, ULK1, and ACC. (**G**) *In vitro* kinase assays using [γ-^32^P]ATP, Purified AMPK, and GST-TFEB fragments purified from bacteria as described and analyzed by SDS-PAGE followed by Coomassie staining and autoradiography. Data are representative of three independent experiments. (**H**) *In vitro* kinase assays using [γ-^32^P]ATP, Purified AMPK, and GST-TFEB fragments purified from bacteria as described and analyzed by SDS-PAGE followed by Coomassie staining and autoradiography. Data are representative of three independent experiments. (**I**) In vitro kinase assays using [γ-32P]ATP, Purified AMPK, and GST-TFE3 fragments purified from bacteria as described and analyzed by SDS-PAGE followed by Coomassie staining and autoradiography. Data are representative of three independent experiments. (**J-K**) *In vitro* kinase assay using cold ATP, Purified AMPK, and GST-TFEB or GST-TFE3 fragments purified from bacteria and analyzed by immunoblotting with the phospho-specific TFEB/TFE3 antibody. (**L**) Endogenous phosphorylated TFEB and TFE3 were immunoprecipitated with pTFEB/pTFE3 antibody followed by immunoblotting using either total TFEB or total TFE3 antibodies. HEK293T WT or AMPK DKO cells were incubated in complete media (Control), starved (EBSS), or in presence of AICAR (2mM) for 2 h. Data are representative of three independent experiments. (**M**) Quantification of the fold change in pTFEB/pTFE3 purified compared to TFEB/TFE3 in total cell lysates (TCL) ratio upon treatment as described in (L) (mean ± SEM of three independent experiments, two-way anova, *P < 0.05; **P < 0.01).

### Phosphorylation of the serine cluster (S466/S467/S469) is required for TFEB activity

To address the biological importance of the potential AMPK-mediated TFEB phosphorylation, we generated MEF cell lines stably expressing either full-length WT TFEB-GFP-3xFLAG or mutant TFEB-S466A-S467A-S469A-GFP-3xFLAG. Expression levels of WT and mutant TFEB proteins were comparable (Fig. 5A). Using a phospho-specific antibody for TFEB Ser211, the major mTORC1 phosphorylation site, we observed a reduction in pS211-TFEB levels in both the WT and mutant upon AICAR and Torin1 treatment demonstrating TFEB dephosphorylation, which should lead to nuclear translocation of WT and mutant TFEB (Fig. 5A). As expected, treatment with both compounds led to decreased mTORC1 activity as measured by immunoblot (Fig. 5A). Moreover, both AICAR and Torin1 treatment led to nuclear localization of WT and S466/467/469A mutant TFEB (Fig. 5B, quantified in C), suggesting that phosphorylation of this serine cluster 466-469 is not important for TFEB nuclear localization. Strikingly, despite its nuclear localization and similar expression levels (Fig. 5A), we observed no increase in TFEB transcriptional activity with the S466/467/469A mutant, when measured by staining for induction of autophagy with DQ-BSA (Fig. 5D, quantified in E), luciferase assay with a CLEAR reporter construct (Fig. 5F), number of lysosomes (Fig. 5G) or by RT-qPCR of known TFEB/TFE3 target genes, upon treatment with AICAR (Fig. 5H) or Torin1 (Fig. 5I). To confirm that TFE3 is regulated similarly by AMPK, we generated MEFs stably expressing full-length WT TFE3-GFP-3xFLAG or mutant TFE3-S567A-S568A-S570A-GFP-3xFLAG. Both proteins are expressed at similar levels (Fig. S3A). AICAR treatment induced nuclear translocation of both proteins (Fig. S3B). However, TFE3 transcriptional activity was not induced with the S567-568-570A mutant upon AICAR treatment, as measured by DQ-BSA assay (Fig. S3C), and CLEAR luciferase reporter assay (Fig. S3D). These data phenocopy the observations made in the context of AMPK loss (Fig. 2), reinforcing an essential role for the presumed AMPK-mediated phosphorylation of TFEB on Ser466/467/469 and TFE3 on Ser 567/568/570 to promote autophagy.

**Figure 5.**
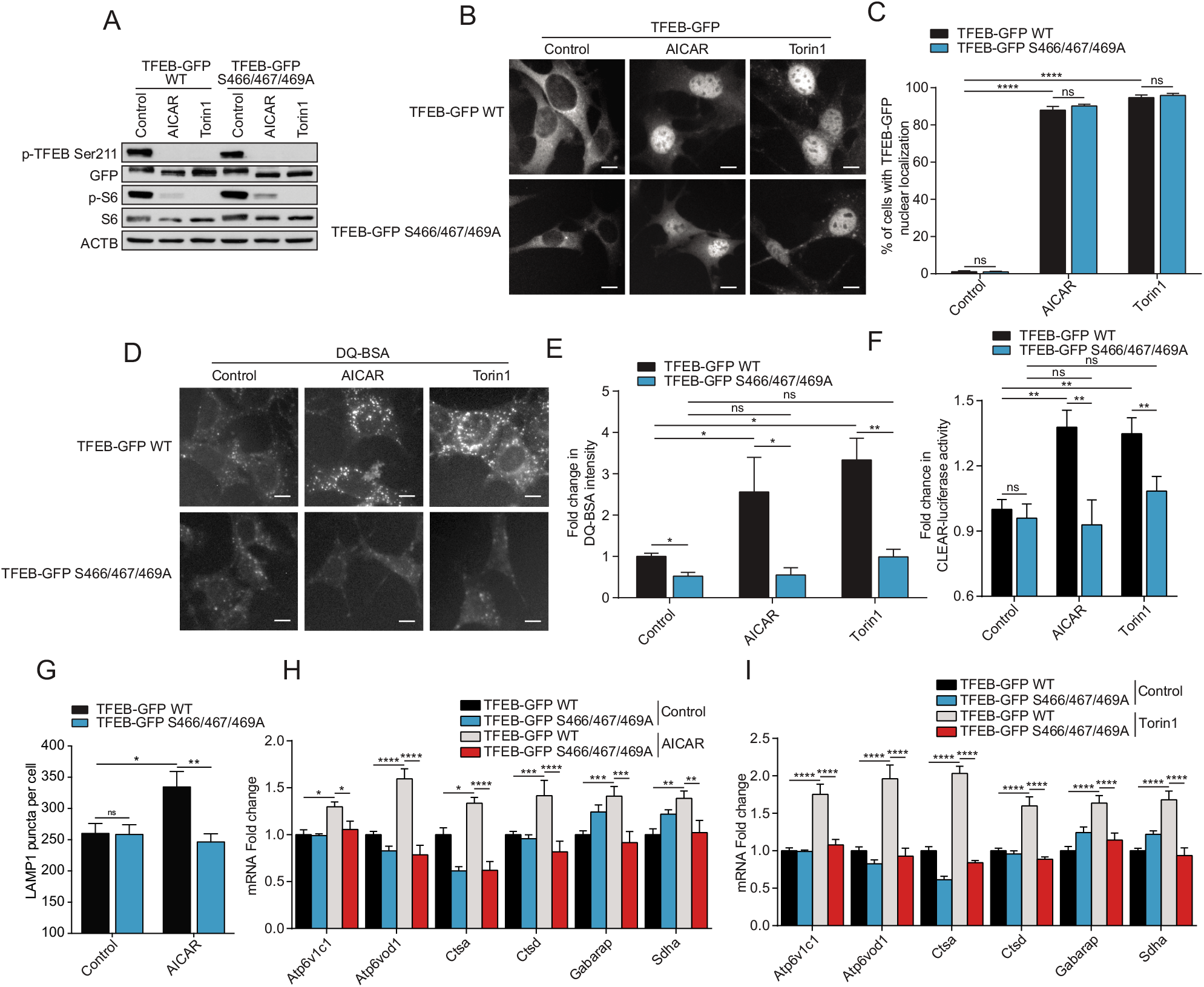
S466/467/469 phosphorylation is required for TFEB activity. (**A**) Immunoblot of protein lysates of MEFs overexpressing TFEB-GFP WT and TFEB-GFP S466/467/469A incubated in complete media (Control) or in presence of AICAR (2 mM) or Torin1 (1 μM) for 2 h. Data are representative of three independent experiments. (**B**) Representative images of MEFs overexpressing TFEB-GFP WT and TFEB-GFP S466/467/469A upon treatment as indicated in (A). Cells were fixed and the GFP fluorescence directly observed by microscopy. Scale bar, 20 μm. Images are representative of three independent experiments. (**C**) Quantification of the percentage of MEFs overexpressing TFEB-GFP WT and TFEB-GFP S466/467/469A with nuclear TFEB-GFP upon treatments as indicated in (A) (mean ± SEM of three independent experiments, two-way ANOVA, ns=not significant, ****P < 0.0001; n > 200 cells per condition). (**D**) Representative images of MEFs expressing TFEB-GFP WT and TFEB-GFP S466/467/469A after 1 h of incubation with DQ-BSA-Red followed by a 2 h chase in complete media in presence of DMSO (Control), AICAR (2 mM) or Torin1 (1 μM) prior to fixation. Scale bar, 20 μm. Images are representative of four independent experiments. (**E**) Relative lysosomal activity, as determined by DQ-BSA assay, in MEFs overexpressing TFEB-GFP WT and TFEB-GFP S466/467/469A upon treatment as indicated in (D) (mean ± SEM of four independent experiments, two-way ANOVA, ns=not significant, *P < 0.05, **P < 0.01; n > 200 cells per condition). (**F**) Relative TFEB/TFE3 transcriptional activity, as determined by CLEAR-luciferase promoter activity normalized against CMV-Renilla in MEFs expressing TFEB-GFP WT and TFEB-GFP S466/467/469A upon treatment as indicated in (A) (mean ± SEM of the luminescence fold change from four independent experiments, two-way ANOVA, ns=not significant, **P < 0.01). (**G**) Quantification of the number of LAMP1 puncta in MEFs overexpressing TFEB-GFP WT and TFEB-GFP S466/467/469A upon treatment as indicated in (A) (mean ± SEM of three independent experiments, two-way ANOVA, ns=not significant, *P < 0.05; **P < 0.01; n > 50 cells per condition). (**H-I**) Relative quantitative real-time PCR analysis of TFEB/TFE3 target genes mRNA transcript levels in MEFs expressing TFEB-GFP WT and TFEB-GFP S466/467/469A upon treatment as indicated in (A) (mean ± SEM of the RNA fold change of indicated mRNAs from four independent experiments, two-way ANOVA, ns=not significant, *P < 0.05, **P< 0.01, ***P < 0.001, ****P < 0.0001).

### S466/467/469A blocks mTORC1-regulated TFEB activity

Next, we investigated whether the mTORC1 and the likely AMPK phosphorylation sites on TFEB affect its activity by using a constitutively active (CA) TFEB mutant. We generated stable MEF lines expressing TFEB-GFP S142A/S211A (TFEB-GFP CA) mutants (mTORC1 phosphorylation sites important for TFEB nuclear translocation) in combination with S466/467/469A mutants (presumed AMPK phosphorylation sites) (Fig. 6A). Expression of the CA mutant resulted in a strong nuclear localization, as expected, which was not affected by the additional S466/467/469A mutations (Fig. 6B, quantified in C). Strikingly, the increased transcriptional activity of the TFEB-GFP CA mutant was abolished upon mutation of the serine residues 466-469, measured by induction of lysosomal activity with DQ-BSA (Fig. 6D, quantified in E), number of LAMP1 puncta (Fig. 6F), CLEAR-luciferase reporter assay (Fig. 6G), and RT-qPCR of TFEB/TFE3 target genes (Fig. 6H).

**Figure 6.**
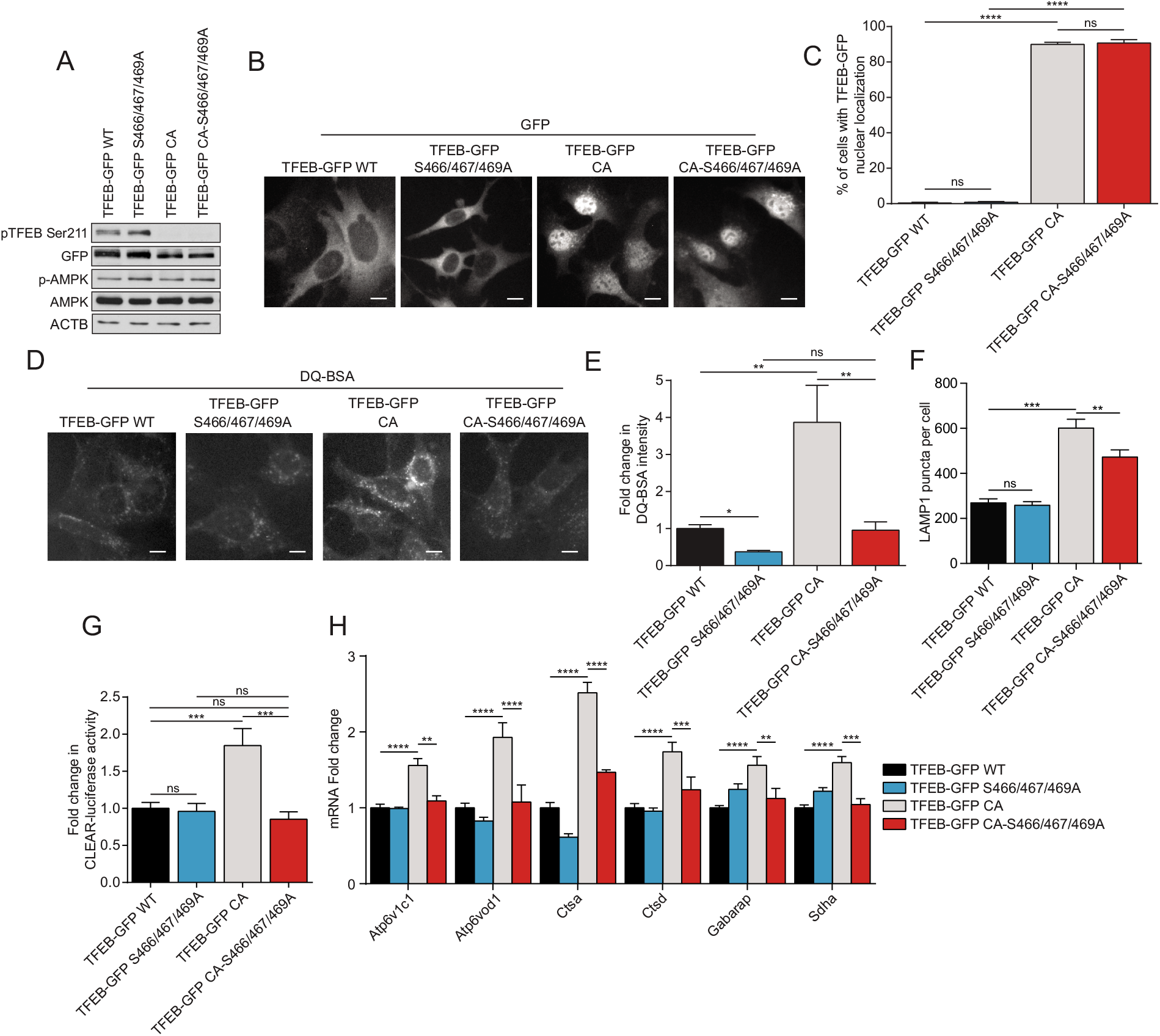
S466/467/469A blocks mTORC1-regulated TFEB activity. (**A**) Immunoblot of protein lysates of MEFs expressing TFEB-GFP WT, TFEB-GFP S466/467/469A, TFEB-GFP S142A/S211A (CA) or TFEB-GFP CA-S466/467/469A incubated in complete media. Data are representative of three independent experiments. (**B**) Representative images of indicated MEF lines incubated in complete media. Cells were fixed and the GFP fluorescence directly observed by microscopy. Scale bar, 20 μm. Images are representative of three independent experiments. (**C**) Quantification of the percentage of MEFs with nuclear TFEB-GFP in complete media (mean ± SEM of three independent experiments, one-way ANOVA, ns=not significant, ****P < 0.0001; n > 200 cells per condition). (**D**) Representative images of MEFs after 1 h of incubation with DQ-BSA-Red followed by a 2 h chase in complete media prior to fixation. Scale bar, 20 μm. Images are representative of four independent experiments. (**E**) Relative lysosomal activity, as determined by DQ-BSA assay, in MEFs upon treatment as indicated in (D) (mean ± SEM of four independent experiments, one-way ANOVA, ns=not significant, **P < 0.01; n > 200 cells per condition). (**F**) Quantification of the number of LAMP1 puncta in MEFs overexpressing TFEB-GFP WT, TFEB-GFP S466/467/469A, TFEB-GFP S142A/S211A (CA) or TFEB-GFP CA+S466/467/469A incubated in complete media (mean ± SEM of three independent experiments, two-way ANOVA, ns=not significant, **P < 0.01; ***P < 0.001; n > 50 cells per condition). (**G**) Relative TFEB/TFE3 transcriptional activity, as determined by CLEARluciferase promoter activity normalized against CMV-Renilla in MEFs were incubated in complete media (mean ± SEM of the luminescence fold change from five independent experiments, one-way ANOVA, ns=not significant, ***P < 0.001). (**H**) Relative quantitative real-time PCR analysis of TFEB/TFE3 target genes mRNA transcript levels in MEFs were incubated in complete media (mean ± SEM of the RNA fold change of indicated mRNAs from four independent experiments, two-way ANOVA, **P< 0.01, ***P < 0.001, ****P < 0.0001).

### Inhibition of TFEB and AMPK sensitized cells to chemotherapy

To further substantiate the physiological relevance of our results with respect to cancer therapy, we measured cell survival following treatment with a chemotherapeutic drug (doxorubicin) in MEFs. First, we tested whether expression of the TFEB gain-of-function mutant (CA-TFEB) results in resistance to chemotherapy. Overexpression of the CA-TFEB mutant, but not wild-type TFEB, conferred increased colony forming ability and resistance to 0.5 nM doxorubicin for 7 days by 108%, which was abolished by mutation of the S466/467/469A sites (Fig. 7A, quantified in B). These observations prompted us to target this drug resistance mechanism pharmacologically. Since no inhibitor of TFEB activity is available, we tested whether inhibition of AMPK, which in turn would block TFEB transcriptional activity, could abolish drug resistance. To this end, we used and validated the specific AMPK inhibitor SBI-0206965. SBI-0206965 was initially identified as an ULK1 inhibitor but was later shown to be more specific toward AMPK [66]. First, we observed that treatment with doxorubicin led to potent activation of AMPK and TFEB, which was reduced upon addition of SBI-0206965 (Fig. 7C-F). Doxorubicin, at a concentration of 0.25 nM for 7 days, did not reduce viability of WT cells (Fig. 7G, quantified in H). However, the combination with SBI-0206965 significantly sensitized the cells to doxorubicin and decreased their colony forming ability (Fig. 7G, quantified in H). At 1.5 μM, SBI-0206965 reduced the percentage of viable cells by 92%. As this AMPK inhibitor is not a clinical grade inhibitor and did not completely blocke AMPK activation, further work would be required to identify more potent compounds for validation studies in animals and humans.

**Figure 7.**
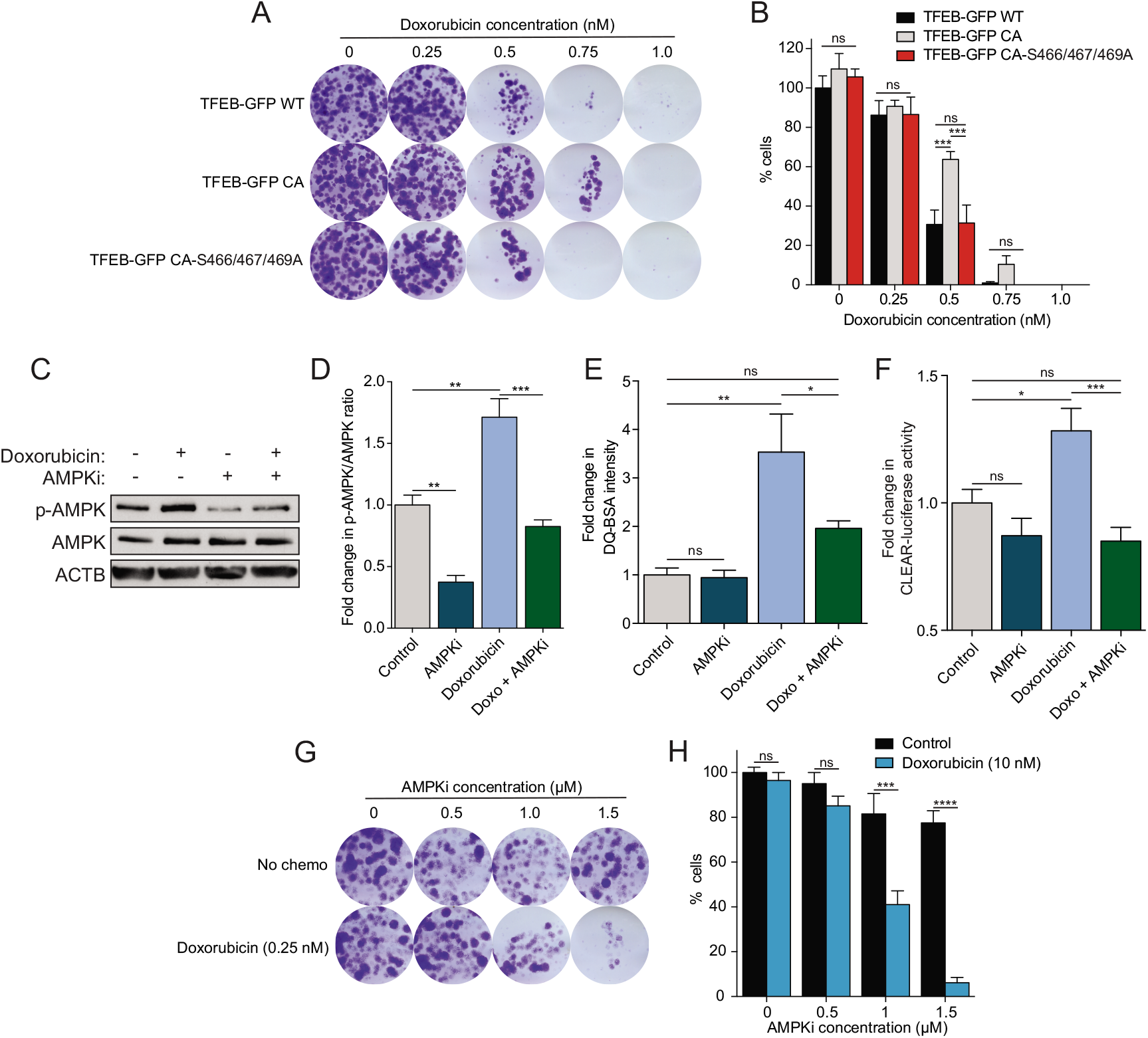
S466/467/469A and AMPK inhibition reduce doxorubicin resistance. (**A**) Representative images of MEFs incubated for 7 days in complete media in presence of indicated doxorubicin concentrations, fixed and stained with crystal violet. Images are representative of three independent experiments. (**B**) Quantification of the percentage of cells remaining upon treatments as indicated in (A) (mean ± SEM of three independent experiments, two-way ANOVA, ns=not significant, ***P <0.001). (**C**) Immunoblot of protein lysates of MEFs WT incubated in complete media in presence of doxorubicin (10 nM) and/or SBI-0206965 (AMPKi, 5 μM) for 24 h. Data are representative of three independent experiments. (**D**) Quantification of the p-AMPK/AMPK ratio upon treatments as indicated in (C) (mean ± SEM of three independent experiments, one-way ANOVA, **P < 0.01, ***P <0.001). (**E**) Relative lysosomal activity, as determined by DQ-BSA assay, in MEFs upon treatment as indicated in (C) (mean ± SEM of four independent experiments, one-way ANOVA, ns=not significant, **P < 0.01; n > 200 cells per condition). (**F**) Relative TFEB/TFE3 transcriptional activity, as determined by CLEAR-luciferase promoter activity normalized against CMV-Renilla in MEFs upon treatment as indicated in (C) (mean ± SEM of the luminescence fold change from five independent experiments, one-way ANOVA, ns=not significant, ***P < 0.001). (**G**) Representative images of MEFs incubated for 7 days in complete medßia in presence of doxorubicin (0.25 nM) and indicated SBI-0206965 (AMPKi) concentrations, fixed and stained with crystal violet. Images are representative of three independent experiments. (**H**) Quantification of the percentage of cells remaining upon treatments as indicated in (G) (mean ± SEM of three independent experiments, two-way ANOVA, ns=not significant, ***P < 0.001, ****P < 0.0001).

Collectively, our data demonstrate that AMPK activation regulates TFEB/TFE3 via two separate mechanisms: a) inhibition of mTORC1 activity that subsequently promotes TFEB/TFE3 nuclear translocation and b) stimulates transcriptional activation. Importantly, we show that AMPK phosphorylates TFEB and TFE3 on specific serine residues and confirm that these residues are essential for their activity. Finally, we show that chemotherapy activates AMPK and TFEB leading to drug resistance, which can be blocked by inhibition of AMPK.

## Discussion

We have previously shown that AMPK and FLCN act upstream of TFEB/TFE3 in regulation of the innate immune response in *C. elegans* and mammalian cells [67]. We reported that FLCN loss constitutively activates AMPK and induces TFEB/TFE3-dependent antimicrobial genes expression. Similar results were obtained upon expression of a constitutively active AMPK mutant in nematodes or pharmacological AMPK activation in mammalian cells. In addition, LPS treatment of macrophages led to acute energy stress, AMPK mediated-TFEB/TFE3 activation, which resulted in increased pro-inflammatory cytokine secretion and phagocytosis. Ablation of AMPK abolished TFEB/TFE3 translocation and activation [67]. These genetic data support the idea that AMPK lies upstream of TFEB/TFE3 and that AMPK regulation of TFEB/TFE3 is necessary for proper autophagy. Here, we report evidence supporting a novel mechanism of TFEB/TFE3 regulation through phosphorylation of TFEB/TFE3 by AMPK (Fig. S4).

In line with previous studies, we showed that severe nutrient starvation such as EBSS starvation (low glucose, no amino acids, no FBS) resulted in both AMPK activation and mTORC1 inhibition [68,69]. The energy drop resulting from the lack of nutrients is acutely sensed by AMPK, which quickly inhibits mTORC1 to stop anabolic processes such as protein synthesis and increase catabolic processes such as autophagy and lysosomal biogenesis. AMPK inhibits mTORC1 via direct phosphorylation of TSC1/TSC2 or by direct phosphorylation of Raptor [36,37]. Our results showed that mTORC1 was inhibited upon 2 h EBSS starvation in both WT and AMPK DKO cells. This confirms that mTORC1 senses the lack of nutrients such as amino acids independently of AMPK when cells are chronically energy starved. mTORC1 senses the presence of amino acids through a multi-protein complex including the Ras-related GTPase (RagA, RagB, RagC, and RagD) guanine nucleotide status (GTP or GDP loaded) [2,50,70,71]. Activated Rags are required to anchor mTORC1 at the lysosomes and phosphorylate TFEB/TFE3 [70]. Rag activation status is regulated by several guanine-exchange factors (GEF) and GTP-ase activating proteins (GAP), particularly by the Ragulator complex, which uses the vacuolar H^+^-adenosine triphosphatase ATPase (v-ATPase) complex to detect the abundance of amino acids [50,54,55,72,73]. In this report, while inactivation of mTORC1 upon EBSS treatment was sufficient to promote TFEB/TFE3 localization to the nucleus independently of AMPK, TFEB/TFE3 activity was abrogated in AMPK DKO cells, suggesting that AMPK dictates the TFEB/TFE3 transcriptional activation status in the nucleus independently of mTORC1. Importantly, the mTORC1 inhibitor Torin1 promoted AMPK-independent nuclear translocation of TFEB/TFE3 but required the presence of AMPK for their transcriptional activation. Despite a slight increase in AMPK activation upon Torin1 treatment, our results suggest that the basal level of AMPK activity is sufficient to promote TFEB/TFE3 activity. However, the kinetics of AMPK phosphorylation remain unclear. Further work would be required to determine if the phosphorylation by AMPK happens at the lysosome or in the nucleus, and if the phosphorylation by mTORC1 precedes those of AMPK.

Consistent with our results, AMPK governs lineage specification by promoting autophagy through transcriptional mechanisms including TFEB [32]. Embryonic stem cells that were deficient for AMPK failed to properly differentiate and had highly deregulated expression of lysosomal genes. Active AMPK resulting from differentiation removed the inhibitory regulation on TFEB by inhibiting mTORC1 downstream of AMPK. Furthermore, AMPK is required to support non-small-cell lung cancer growth by regulating lysosomal gene expression via TFE3 [74]. Upon glucose starvation, AMPK activity is required to promote TFE3 nuclear localization and transcriptional activity. Our results are consistent with previous studies, but we add another layer of complexity in TFEB regulation. AMPK was required not only to inhibit mTORC1 to promote TFEB/TFE3 nuclear localization, but also to stimulate TFEB/TFE3 transcriptional activity upon acute energy stress.

The only reported TFEB transcriptional coactivator is the Coactivator-associated arginine methyltransferase 1 (CARM1), which is induced in an AMPK-dependent manner upon chronic glucose starvation for 12-24 h [33]. Chronic glucose starvation promoted AMPK-dependent Forkhead box O3 (FOXO3) phosphorylation in the nucleus, which transcriptionally repressed the E3 ubiquitin ligase component S-phase kinase-associated protein 2 (SKP2) leading to increased levels of CARM1. The kinetics of starvation and activation observed in this study were different from our study; increased levels of CARM1 protein and reduction of SKP2 required 12-24 h of glucose starvation. We observed an increase in TFEB/TFE3 activity after 15 minutes of EBSS starvation with a peak after 2 h. Moreover, in this study SKP2 silencing in AMPK DKO MEFs only partially restored autophagic activity, suggesting that AMPK also regulates TFEB/TFE3 activity via other pathways. This is consistent with a distinct response upon acute versus chronic starvation stress; AMPK quickly promotes autophagy via direct phosphorylation of ULK1 and TFEB/TFE3 and inhibition of mTORC1, while long-term starvation might further promote autophagy epigenetically by transcriptionally modulating TFEB/TFE3 co-activators, such as CARM1.

TFEB subcellular localization and activity is regulated by phosphorylation through multiple kinases. We report that AMPK may activate TFEB via phosphorylation on a highly conserved cluster of serine residues near the C-terminus. Mutation of S466/467/469A led to abrogation of TFEB activation upon AMPK activation or mTORC1 inhibition. It is conceivable that a consequence of AMPK-mediated phosphorylation of S466/467/469 is a conformational change that renders TFEB transcriptionally active. Further work is required to delineate the mechanism through which mutation of this region reduces TFEB transcriptional activity. One possibility is that these AMPK serine phosphorylation sites control the interaction of TFEB/TFE3 with yet unknown transcriptional coactivators.

Interestingly, Protein kinase B (PKB, AKT) is reported to phosphorylate TFEB at serine 467, but this phosphorylation led to repression of TFEB nuclear translocation independently of mTORC1 [75]. In this study, mutation of S467 to alanine led to increased TFEB nuclear localization. In our hands this mutant S467A had no effect on TFEB nuclear translocation. In agreement, the triple mutant S466/467/469A also did not affect nuclear localization. However, the triple-mutant resulted in reduced autophagy, lysosomal activity, CLEAR reporter activity and TFEB/TFE3 target gene expression. AMPK and PKB might both phosphorylate the C-terminal region in a different hierarchy which could result in different activity. Similar regulation by AMPK and PKB of a different transcription factor was recently described [76]. Consistent with our findings, it is reported that a TFEB mutant harboring a C-terminal truncated of 15 residues, which contains the S466/467/469 AMPK phosphorylation sites, lacked transcriptional activity [77]. Phosphorylation of S462, 463, 467, and 469 is shown to be important for the stability and activity of TFEB [77]. In our study, we showed that phosphorylation of the C-terminal region of TFEB by AMPK promoted TFEB transcriptional activity, but we did not observe an effect on protein levels upon mutation of S466/467/469A compared to WT. The C-terminal region of TFEB/TFE3 might be tightly regulated via phosphorylation through multiple kinases; the phosphorylation status of the many serine residues located at this region could either promote or inhibit TFEB/TFE3 activity depending on the phosphorylation pattern.

We were not able to determine how many of the three serines were phosphorylated in cultured cells from our mass spectrometry experiments. However, it was necessary to mutate all three serines residues for full abrogation of transcriptional activity in cells. Single or double combination mutations of the three serines led to partial responses. It is likely that AMPK phosphorylates the adjacent serine(s) if only one or two of the three serines is mutated to alanine. Moreover, we could not fully demonstrate that AMPK phosphorylates all three sites of TFEB in cells. However, we developed a phospho-specific antibody against a peptide containing all three phospho-serines of TFEB and TFE3 and were able to validate this antibody *in vitro* and in cultured cells. More work involving phosphospecific antibodies and *in vivo* metabolic labelling is required to determine the exact site(s) of phosphorylation.

Several chromosomal translocations of *TFEB/TFE3* with strong promoters, such as *MALAT1, PRCC, ASPSCR1, SFPQ, NONO* and *CLTC*, have been strongly linked with increased TFEB/TFE3 activity, renal cell carcinoma (RCC) and alveolar soft part sarcoma [6,14,15]. Knock-down of the MiT/TFE family in pancreatic ductal adenocarcinoma (PDA) cells reduced their growth in xenograft assays, while overexpression increased their survival in response to nutrient stress and promoted tumorigenesis in orthotopic injection [78,79]. In addition to increased proliferation, TFEB/TFE3 are linked to chemotherapy resistance through lysosomal sequestration. Several anticancer drugs, such as doxorubicin, have been shown to be trapped in lysosomes due to their hydrophobic weak base properties [80,81]. Therefore, stresses such as nutrient starvation and chemotherapy treatment, augment lysosomal biogenesis, increase the number and size of lysosomes per cell, enhance drug sequestration and thus, confer drug resistance [82,83]. In our work, we reported that AMPK-mediated phosphorylation of TFEB was required for chemotherapy resistance. Moreover, the AMPK inhibitor SBI-0206965 sensitized the cells to doxorubicin, suggesting that inhibiting AMPK might be a potential target to reduce chemotherapy resistance. Our study represents a proof of concept and would benefit from the development of more potent inhibitors of AMPK.

Finally, it was reported that AMPK also regulates autophagy via non-transcriptional regulation such as direct phosphorylation of the autophagy initiating kinase ULK1 [29,84]. Our data reveal strong similarities between the dual and opposing regulations of ULK1 and TFEB/TFE3 by AMPK and mTORC1. AMPK directly phosphorylates and activates ULK1 (on serines S317, S467, S555, S574, S637, and S777), while mTORC1 inhibits ULK1 by directly phosphorylating serine residue S757. Together, our present and previous results demonstrate that acute nutrient starvation leads to a) AMPK-mediated inhibition of mTORC1 that promotes ULK1 dephosphorylation at S757 and TFEB dephosphorylation at S142 and S211, which are necessary steps for activation, and b) AMPK-mediated phosphorylation of ULK1 and TFEB at multiple sites, which complete the activation of both proteins. Both, AMPK-mediated phosphorylation of ULK1 and TFEB and inhibition of mTORC1-dependent dephosphorylation of TFEB and ULK1 are required for activity of these autophagy regulatory proteins. Strikingly, AMPK and mTORC1 regulate both critical regulators of autophagy ULK1 and TFEB/TFE3 in a similar manner. In conclusion, these results suggest that AMPK activates autophagy via a dual ‘fail-safe’ mechanism involving not only inactivation of mTORC1 but also phosphorylation of ULK1 and TFEB/TFE3.

## Material and Methods

### Cell lines culture and treatments

Primary MEFs were isolated from C57BL/6 E12.5 *Flcn* floxed mice (generously provided by Dr. L.S. Schmidt, NCI, Bethesda, MD, USA) or *Flcn/Prkaa1/Prkaa2* floxed mice and cultured as described previously [41]. FLCN and FLCN/AMPK KO MEFs were generated after immortalization of primary Flox/Flox MEFs with SV40 large T and retroviral infection with a Cre recombinase, followed by puromycin selection (Bioshop Canada, PUR333.25). WT and AMPK DKO MEFs cells were generously provided by Dr. Benoit Viollet (Institut Cochin INSERM, Paris, France). HEK293T AMPK DKO cells were generated by co-transfection of lentivirus containing CRISPR-Cas9 encoding plasmid and target gRNA of both AMPK subunits *(PRKAA1;* gRNA sequence GAAGATCGGCCACTACATTC, and *PRKAA2;* gRNA sequence AAGATCGGACACTACGTGCT) and single-cell cloned. HEK293T control cell lines were also single-cell cloned after transduction with non-coding gRNA and CRISPR-Cas9. Cell lines were maintained in Dulbecco’s modified Eagle’s medium (DMEM) (Wisent, 319-005CL) supplemented with 10% fetal bovine serum (FBS) (Wisent, 080-150), 100 U/ml penicillin+100 μg/ml streptomycin (Wisent, 450-201-EL), and 50 μg/mL gentamycin (Wisent, 450-135) in 5% CO2 at 37°C. MEFs stably overexpressing TFEB-GFP-3xFLAG were transduced with retrovirus, selected with zeocin and single-cell cloned. MEFs stably expressing inducible Poly94Q-CFP chains were transduced with lentivirus and selected with hygromycin B (Thermo Fisher Scientific, 10687010). Polyglutamine aggregates were induced by addition of Doxycyclin (5 μg/mL, Sigma-Aldrich, D9891) for 3 days to culture media prior to drug treatments. For EBSS starvation experiments, cells were washed twice with PBS and incubated in Earle’s Balanced Salt Solution (EBSS) (Sigma-Aldrich, E2888) for 2 h. For drug treatment experiments, cells were incubated for 2 h in medium containing one of the following reagents: Dimethyl sulfoxide (DMSO) (0.1%) (Bioshop Canada, DMS666), Torin1 (1 μM, Tocris, 10-1-4247), and AICAR (2 mM, Enzo Life Sciences, 89158-090). For chemotherapy drug resistance experiments, cells were incubated for either 24 h or 7 days in medium containing one or a combination of the following reagents: DMSO (Bioshop Canada, DMS666), Doxorubicin (Sigma-Aldrich, D1515), and SBI-0206965 (APExBIO, A8715-5). The final DMSO concentration never exceeded 0.1% and this concentration was shown to have no detrimental effect on all the studied cells.

### Antibodies

The FLCN rabbit polyclonal antibody was generated by the McGill animal resource centre using recombinant GST-FLCN. The p-TFEB S211 and p-TFEB S466/467/469 / p-TFE3 S567/568/570 antibodies were generated by Yenzym Antibodies, LLC. TFEB (Bethyl Laboratories, A303-673A), TFE3 (Cell Signaling Technology, 14779S), AMPKα (Cell Signaling Technology, 2532), p-AMPKα (Thr172) (Cell Signaling Technology, 2531), ACC1/2 (Cell Signaling Technology, 3676), p-ACC1/2 (S79) (Cell Signaling Technology, 3661), 4EBP1 (Cell Signaling Technology, 9644), p-4EBP1 (Cell Signaling Technology, 9456), S6 (Cell Signaling Technology, 2217), p-S6 (Cell Signaling Technology, 4858), LaminA (Santa Cruz Biotechnology, sc-71481), ß-Actin (Santa Cruz Biotechnology, sc-47778), Tubulin (Sigma-Aldrich, T9026), Anti-Polyglutamine-Expansion (Millipore, MAB1574), anti-GFP (Clontech, 632460), LAMP1 (Abcam, ab24170) antibodies are commercially available.

### Immunofluorescence

Cells were gently washed with PBS and fixed in petri dishes with 3.7% formaldehyde (Bioshop Canada, FOR201.500) at room temperature for 30 min. After fixation, cells were washed twice with PBS and then permeabilized with 0.3% Triton X-100 (Bioshop Canada, TRX506.100) in PBS at room temperature for 10 min. Cells were incubated in 5% BSA (Bioshop Canada, ALB001.500) in PBS for 1 h and then with TFEB or TFE3 primary antibodies in 5% BSA in PBS for 2.5 h at 37°C. Cells were washed three times with PBS and incubated with the corresponding secondary antibodies conjugated to Alexa Fluor 488 (Thermo Fisher Scientific, A-11008) and DAPI (2 μg/ml) (Molecular Probes, D-1306) in 5% BSA in PBS for 30 min at 37°C. PBS-washed dishes were covered with cover slips and observed with Axioskop microscope (Zeiss). Scoring of the percentage of cells displaying nuclear localization was done manually using at least three different frames containing at least 100 cells.

### Lysosome number and size

Cells were grown in 6-wells on coverslips. After treatment as described above, cells were gently washed twice with PBS and fixed with 4% paraformaldehyde (Bioshop Canada, par070.500) at room temperature for 15 min. After fixation, cells were washed twice with PBS, permeabilized with 0.1% saponin in PBS at room temperature for 10 min. Cells were incubated with LAMP1 primary antibody in 0.1% saponin (Alfa Aesar, J63209-AK), 1% FBS in PBS for 2 h in a humidity chamber at 37°C. Cells were washed three times with PBS and incubated with Alexa Fluor 594 Anti-rabbit (Thermo Fisher, A32740), Alexa Fluor 647 Phalloidin (Thermo Fisher, A22287), and DAPI (2 μg/ml) in 0.1% saponin, 1% FBS in PBS for 1 h in a humidity chamber at 37°C. PBS-washed coverslips were mounted on microscope slides and observed on confocal microscope LSM-800 (Zeiss) with 64x oil-immersion objectives at a zoom of 0.8 and acquired using LSM software. Lysosome number and size were quantified using Metamorph (Molecular Devices, LLC.) from at least 50 individual cells per condition.

### DQ-BSA assay

MEFs were incubated with 5 μg/mL DQ Red BSA (Thermo Fisher Scientific, D12051) for 1 h and washed twice with 37°C PBS. EBSS or DMEM containing AICAR, Torin1, or vehicle were added for an additional 2 h. Cells were then fixed and stained for TFEB or TFE3 and DAPI as described above. Grey pixels from pictures acquired on Axioskop microscope (Zeiss) were quantified using ImageJ (NIH).

### Protein extraction and immunoblotting

Cells were washed twice with PBS, lysed in AMPK lysis buffer (10 mM Tris-HCl (pH 8.0), 0.5 mM CHAPS, 1.5 mM MgCl2 1 mM EGTA, 10% glycerol, 5 mM NaF, 0.1 mM Sodium orthovanadate (Na3VO4), 1 mM benzamidine, 5 mM Sodium pyrophosphate (NaPPi), supplemented with a complete protease inhibitor cocktail (Roche, 5056489001) and Dithiothreitol (DTT) (1 mM), sonicated 2 x 30 sec at maximum power, and cell lysates were cleared by centrifugation at 13000 x *g*. Proteins were separated on SDS-PAGE gels and revealed by western blot as previously described [41] using the antibodies listed above, and quantified using ImageJ.

### Htt94Q aggregation assay

Cells stably expressing doxycycline-inducible Htt94Q-CFP fusion plasmids were plated and incubated with 5 μg/mL of doxycycline for 3 days prior to treatment with AICAR (2 mM), Torin1 (1 μM), or vehicle for 2 h. Cells were fixed in petri dishes with 3.7% formaldehyde at room temperature for 30 min. pTreTight-Htt94Q-CFP was a gift from Nico Dantuma (Addgene plasmid # 23966; http://n2t.net/addgene:23966; RRID:Addgene_23966). PBS-washed dishes were covered with cover slips and observed with Axioskop microscope (Zeiss). Scoring of the percentage of cells displaying CFP puncta was done manually using at least three different frames containing at least 200 cells.

For soluble/insoluble Htt94Q-CFP quantification, Cells were lysed in soluble lysis buffer (1% Triton X-100, phosphatase/protease inhibitors in PBS) on ice for 30 min, followed by centrifugation at 15,000 x *g* for 30 min at 4°C. Soluble fraction was collected and the pellets were washed 4 times in soluble lysis buffer. The remaining pellets were solubilized using insoluble lysis buffer (1% SDS, 1% Triton X-100, phosphatase/protease inhibitors in PBS) at 60°C for 1 h. After centrifugation at 15,000 x *g* for 30 min at 4°C, the insoluble fraction was collected. After revelation by western blot as described above, multiple exposures were used to quantify the relative amount of insoluble Htt94Q-CFP using ImageJ (NIH).

### CLEAR-luciferase reporter assays

MEFs were seeded in 6-well plates and transfected for 6-8 h with 1 μg of plasmid 4XCLEAR-luciferase reporter (Addgene, 66800) and 0.1 ug of CMV-Renilla Luciferase (Promega, E2261) using 5 μL of Polyethylenimine (PEI) (Polysciences, 23966-1) at 1 mg/mL stock concentration. 24 h after transfection, cells were treated with AICAR, Torin1, EBSS, or vehicle for 2 h. Proteins were extracted using 100 μL of Passive Lysis Buffer from Dual-Luciferase reporter assay systems (Promega, E1980) according to manufacturer’s instruction and assayed using FLUOstar Omega (BMG Labtech). Samples were normalized against non-transfected controls and CMV-Renilla values.

### Quantitative real-time PCR

After treatments described above, cells were collected, and total RNA was isolated and purified using Total RNA Mini Kit (Geneaid, RB300) according to the manufacturer’s instructions. For quantitative real-time PCR analysis, 0.75 μg of total RNA was reverse-transcribed using the iScript Reverse Transcription Supermix for RT-qPCR (BioRad, 1708841). SYBR Green reactions using the SYBR Green qPCR supermix (BioRad, 1725125) and specific primers (Table 1) were performed using an AriaMX Real-time PCR system (Agilent Technologies). Relative expression of mRNAs was determined after normalization against housekeeping genes TBP, B2M, or 18S.

### Co-immunoprecipitations

HEK293T cells were lysed in lysis buffer (25 mM Tris, pH 7.4, 150 mM NaCl, 5 mM MgCl2, 0.5% NP-40, 1 mM Dithiothreitol (DTT), 5% glycerol, 1 mM phenylmethanesulfonyl fluoride, 2 μg/ml pepstatin, 2 μg/ml leupeptin, and 2 μg/ml aprotinin) in the presence of phosphatase-inhibitors (Sigma-Aldrich, 4906837001) after one wash with cold PBS. Cellular debris was removed by centrifugation at 13,000 x *g* for 10 min at 4°C and the lysate pre-cleared for 30 min with G-agarose beads. The cell lysates were immunoprecipitated for 2 h at 4°C, using and anti-AMPK antibodies (2 μg) or p-TFEB/p-TFE3 antibody (2 μg). 50 % Protein G-agarose beads (30 μl) (Millipore, 16-266) were added to lysates after 1 h. Beads were washed once with lysis buffer for 5 minutes, followed by 2 washes with wash buffer (25 mM Tris, pH 7.4, 150 mM NaCl, 5 mM MgCl2, 1 mM Dithiothreitol (DTT), 5% glycerol, 1 mM phenylmethanesulfonyl fluoride, 2 μg/ml pepstatin, 2 μg/ml leupeptin, and 2 μg/ml aprotinin). SDS-PAGE buffer was added to the washed beads and half of the volume was separated on a 4 – 15 % polyacrylamide, precast Mini-PROTEAN TGS gel (BioRad, 4561094) followed by transfer on Trans-Blot Turbo Transfect Pack (0.2 μm) nitrocellulose (BioRad, 1620115), and western blotting was performed as described with indicated antibodies.

### Protein purification from cells

HEK293T were seeded in 150 mm plates and transfected for 8 h with 10 μg of a TFEB-GFP-3xFLAG plasmid using 100 mM Polyethylenimine (PEI). After 15 min on ice, debris was cleared by centrifugation for 10 min at 13000 x *g* at 4°C. Lysates were pre-cleared on rotator using Protein A + G mix (Millipore, 16-156 and 16-266) for 1 h at 4°C, followed by TFEB immunoprecipitation for 3 h using magnetic Flag beads (Sigma-Aldrich, M8823-5ML). Beads were washed 3 times with lysis buffer and TFEB or TFE3 eluted using 200 ng/μL 3X-flag-peptide (Sigma-Aldrich, F4799-4MG).

### Protein purification from bacteria

TFEB fragments were cloned in frame with GST in pGEX-2T (GE Healthcare, 27-1542-01). Overnight bacterial culture *(E. coli*, DH5α or BL21) was diluted 1:100 and incubated with shaking at 37°C in LB + ampicillin (100 μg/ml) until the OD600 of 0.6. Protein expression was induced by adding 0.5 mM IPTG (Bioshop Canada, IPT002.1) and incubation for an extra 2.5 h at 37°C. Bacteria were pelleted 10 min at 7700 x *g*, re-suspended in ice-cold PBS and sonicated 3 x 10 sec at maximum power. Triton X-100 to a final concentration of 1% was added and mixed at 4°C for 30 min. Debris was pelleted for 10 min at 12000 x *g* and the supernatant incubated with PBS-washed glutathione-sepharose 4FastFlow beads (GE Healthcare, 17075601) overnight at 4°C. The beads were washed in PBS and the proteins eluted by incubating for 15 min with 50 mM Tris-HCl (pH 8.0) and 10 mM glutathione (Bioshop Canada, GTH001.5).

### Kinase assay

Bacteria or cell purified proteins or recombinant TFEB (LD Biopharma, hTF-1724), ULK1 (Sigma-Aldrich, SRP5096-10UG), ACC (Sigma-Aldrich, A6986-10UG) were incubated in presence of [γ-^32^P] ATP (Perkin-Elmer, BLU502H250UC) on ice for 10 min in reaction buffer (25 mM Hepes pH 7.4, 50 mM KCl, 10 mM MgCl2, 100 μM ATP, 150 μM AMP, 5 mM Sodium pyrophosphate (NaPPi), complete protease inhibitor cocktail (Roche, 5056489001). Recombinant active AMPK (Millipore, 14-840) was added and incubated for 30 min at 30°C. Reaction was stopped by adding Laemmli sample buffer and incubation at 95°C for 5 min. Proteins were resolved on acrylamide gels, coomassie stained, dried, and radioactivity was revealed with Typhoon Trio (GE Healthcare).

### Mass spectrometric analysis of TFEB phosphorylations

TFEB fragments were purified, incubated in presence of AMPK in a kinase assay reaction buffer, and sent to PhenoSwitch Bioscience for analysis. TFEB fragments were digested using 8 μl proteomics grade Trypsin in 50 mM ammonium bicarbonate, at a concentration of 12 ng/μl, overnight. The digested protein samples were acidified in 0.1% trifluoroacetic acid (TFA) in 50% acetonitrile (ACN) and subjected to phosphopeptide enrichment, using titanium dioxide beads, without competition. Briefly, peptide digested in 400 μl 50% ACN, 0.1% TFA were incubated for 30 min with 3-4 μl of TiO2 beads (Titansphere, Canadian Biosciences). The beads were pelleted by centrifugation for 30 sec in a table top centrifuge and washed once with 400 μl 50% ACN, 0.1% TFA. After pelleting the TiO2 beads a second time, bound phosphopeptides were eluted with 100 μl 5% ammonium hydroxide in 50% ACN. The eluted phosphopeptide samples were dried in a speed vac. The enriched phosphopeptide samples were dissolved in 25 μl of demineralized water, 0.1% FA. They were subjected to reverse phase separation, on a 15-cm column (Acclaim PepMap RPLC, Thermo Fisher Scientific) using a nano UHPLC (Easy nLC-1000, Thermo Fisher Scientific) running a flow rate of 350 nl per min and a water/ACN gradient spanning 3-35% ACN in 100 min. The eluted peptides were analyzed using a Q-Exactive HF orbitrap mass spectrometer (Thermo Fisher Scientific), operating at 120,000 resolution for precursor scans, in the mass range from 350 to 1500 m/z, with a trap fill set to 5e6 and at 30,000 resolution for MSMS scans in the mass range from 100-2000 m/z, with a trap fill set to 5e4. Dynamic exclusion was set to 8 sec and the instrument was run at 25 MSMS analyses per duty cycle. The resulting .raw files were converted into Mascot Generic Files (mgf) using Peak Distiller (Matrix Sciences) and searched against the Uniprot reviewed human proteome (UP000005640) using Mascot v. 2.5 (Matrix Sciences). Precursor mass tolerance was set at 5 ppm and MSMS mass tolerance was set at 50 mmu. Variable modifications were allowed for oxidized methionine and for phosphorylations on serines, threonines and tyrosines. Data were thereafter re-searched, using X!Tandem and the data were validated and false discovery rates (FDR) were determined, using Scaffold 4 (Proteome Software). Data were filtered, setting the peptide FDR cut-off to 2%.

### Doxorubicin resistance assays

MEFs were incubated with the indicated concentrations of doxorubicin and/or SBI-0206965 (Apexbio, A8715-5) and/or vehicle (DMSO) for 7 days. Viability was assessed in 6-well plates. After 7 days, cells were fixed in petri dishes with 3.7% formaldehyde at room temperature for 60 min. After fixation, cells were incubated 60 min with 0.1% crystal violet solution, thoroughly washed in water and allowed to dry. Quantification was performed using ImageJ software.

### Statistical Analysis

Data are expressed as mean ±SEM. All experiments were performed at least three times as indicated. Statistical analyses for all data were performed using student’s t-test, one-way ANOVA or two-way ANOVA as indicated using GraphPad Prism 7 software. Statistical significance is indicated in figures (*P<0.05, **P<0.01, ***P<0.001, ****P<0.0001).

## Abbreviations

ACC1/2: acetyl-CoA carboxylase 1 or 2
ACTB: actin beta
AICAR: 5-aminoimidazole-4-carboxamide ribonucleotide
AMPK: AMP-activated protein kinase
AMPKi: AMPK inhibitor, SBI-0206965
CA: constitutively active
CARM1: coactivator-associated arginine methyltransferase
CFP: cyan fluorescent protein
CLEAR: coordinated lysosomal expression and regulation
DKO: double knockout
DMEM: Dulbecco’s modified Eagle’s medium
DMSO: dimethyl sulfoxide
DQ-BSA: selfquenched BODIPY^®^dye conjugates of bovine serum albumin
EBSS: Earle’s balanced salt solution
FLCN: folliculin
GFP: green fluorescent protein
GST: glutathione S-transferases
HD: Huntington disease
HTT: huntingtin
KO: knock-out
LAMP1: lysosomal-associated membrane protein 1
MEF: mouse embryonic fibroblasts
MITF: melanocyte inducing transcription factor
mTORC1: mechanistic target of rapamycin complex 1
PolyQ: polyglutamine
RT-qPCR: reverse transcription quantitative polymerase chain reaction
S6: ribosomal protein S6
TCL: total cell lysates
TFE3: transcription factor E3
TFEB: transcription factor EB
TKO: triple knock-out
ULK1: unc-51-like kinase 1

## Acknowledgements

M.P., L.E-H., were supported by FRQS/CIHR, and FRQS, respectively. This work was supported by grants to A.P. from the Kidney Foundation of Canada, Terry Fox foundation (TFF-166128), CIHR (PJT-165829) and the Cancer Research Society. We would like to thank Paula Coehlo, the McGill University Life Sciences Complex Advanced BioImaging Facility (ABIF), and McGill University Mass Spectrometry Facility for their support and advices.

## Author contributions

Conception and design of the experiments: M.P., L.EH., L.C.Z., P.B., and A.P.; collection, assembly, analysis and interpretation of data: M.P., L.EH., L.C.Z., K.D., P.P., H.J., and P.B.; drafting the article or revising it critically for important intellectual content: M.P., L.EH., L.C.Z., P.B., P.M.S. and A.P.

