## Supplemental information for "AMPK-dependent phosphorylation is required for transcriptional activation of TFEB/TFE3"

Supplementary tables

**Table 1.** List of primers used in RT-qPCR experiments.

| Gene | Forward primers | Reverse primers |
|---|---|---|
| **mouse-B2M** | CTGACCGGCCTGTATGCTAT | CCGTTCTTCAGCATTTGGAT |
| **mouse-TBP** | ACCTTATGCTCAGGGCTTGG | GCCATAAGGCATCATTGGAC |
| **mouse-ATPV1C1** | CCAGCGGAGTTGACTTGGTT | ATTGTATGCAGACGCCCGAG |
| **mouse-ATPV0D1** | TCCGCCACATGAGAAACCAT | CTCAAAGCTGCCTAGCGGAT |
| **mouse-CTSA** | GAGCAGAACGACAACTCCCT | TGCCCACAATTCGAGACACT |
| **mouse-CTSD** | CTATAAGCCGGCGACCTCTG | TGAACTTGCGCAGAGGGATT |
| **mouse-GABARAP** | CGTGCTGAAGATGCCTTGTT | AAGGAGCGCCACCTCTCTTC |
| **mouse-SDHA** | TTACCTGCGTTTCCCCTCAT | AAGTCTGGCGCAACTCAATC |
| **human-18S** | AACCCGTTGAACCCCATT | CCATCCAATCGGTAGTAGCG |
| **human-TBP** | AGGGTTTCTGGTTTGCCAAGA | CTGAATAGGCTGTGGGGTCA |
| **human-ATPV1C1** | ATTGCATGCGGCAACTTCAA | CCAAGACATCCAACGTGCCA |
| **human-ATPV0D1** | CCATGTCGTTCTTCCCGGAG | TCAGAGGTGATGCCTCGTTG |
| **human-CTSA** | CCAACACAACAGCTGCTTCC | CTGGGAGTTCATGCTTCGGT |
| **human-CTSD** | TCTGTGGAGGACCTGATTGC | CGATGCCAATCTCCCCGTAG |
| **human-GABARAP** | AGAAGAGCATCCGTTCGAGA | TCTACTATCACCGGCACCCG |
| **human-SDHA** | TGGAGCTGCAGAACCTGATG | CTCCAGTGCTCCTCAAAGGG |

Supplementary figures

Supplementary Figure 1

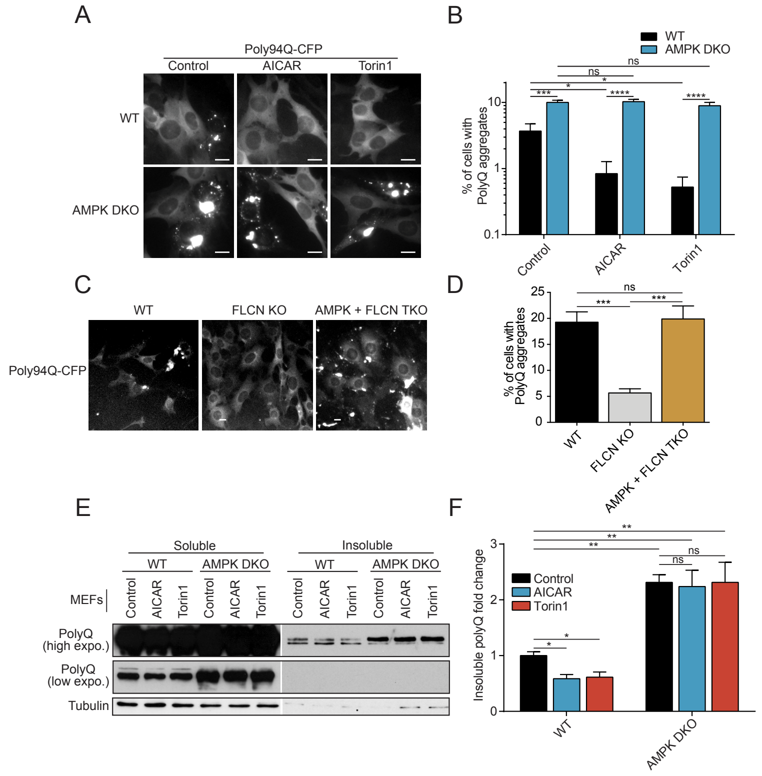


### Figure S1. AMPK is required for protein aggregates clearance in a model of Huntington disease

(**A**) Representative images of MEFs WT and AMPK KO expressing a doxycycline-inducible fusion protein consisting of the Cyan Fluorescent Protein (CFP) fused to a chain of 94 glutamine residues induced for 3 days, followed by incubation in complete media in presence of DMSO (Control), AICAR (2 mM) or Torin1 (1 µM) for 2 h prior to fixation. Scale bar, 20 µm. Data are representative of four independent experiments. (**B**) Quantification of the percentage of MEFs expressing PolyQ-CFP with aggregates accumulation upon treatments as indicated in (A) (mean ± SEM of four independent experiments, two-way

ANOVA versus WT control-treated cells, ns=not significant, *P < 0.05, ***P < 0.001, ****P < 0.0001; n > 200 cells per condition). (**C**) Representative images of MEFs WT, FLCN KO and AMPK/FLCN TKO expressing a doxycycline-inducible fusion protein consisting of the Cyan Fluorescent Protein (CFP) fused to a chain of 94 glutamine residues induced for 3 days prior to fixation. Scale bar, 20 µm. Data are representative of three independent experiments. (**D**) Quantification of the percentage of MEFs expressing PolyQ-CFP with aggregates accumulation (mean ± SEM of three independent experiments, two-way ANOVA, ns=not significant, ***P < 0.001; n > 200 cells per condition). (**E**) Immunoblot of protein lysates resulting from soluble-insoluble subcellular fractionation of MEFs expressing PolyQ-CFP treated as indicated in (A). Data are representative of three independent experiments. (**F**) Quantification of insoluble polyQ protein levels in MEFs expressing PolyQ-CFP treated as indicated in (A) (mean ± SEM of the fold change from three independent experiments, two-way ANOVA, ns=not significant, *P < 0.05, **P < 0.01).

Supplementary Figure 2

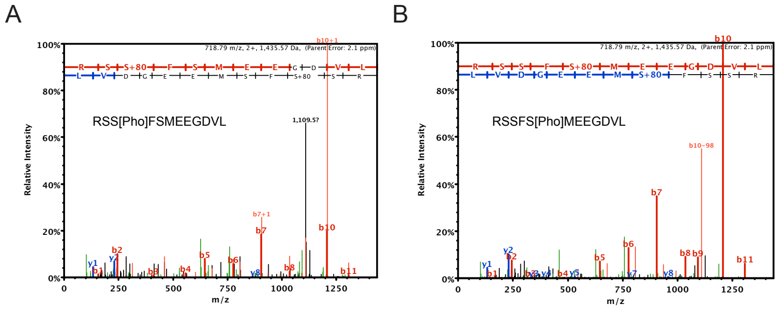


### Figure S2. Mass spectrometry spectra

(**A-B**) MS/MS spectrum of phospho-peptides representing residues 464-476 of TFEB. GST-TFEB C-terminal fragment was purified from bacteria, subjected to in vitro kinase assays with recombinant AMPK and analyzed by mass spectrometry. As shown, collision-induced dissociation spectra of the doubly charged parent ion resulted in several b and y series fragment ions (b and y ions are, respectively, the N- or C-terminal fragments produced when the parent peptide is fragmented at peptide bonds). The spectra shown contain enough sequence information to conclusively localize phosphorylation sites to both S467 (A) and S469 (B).

Supplementary Figure 3

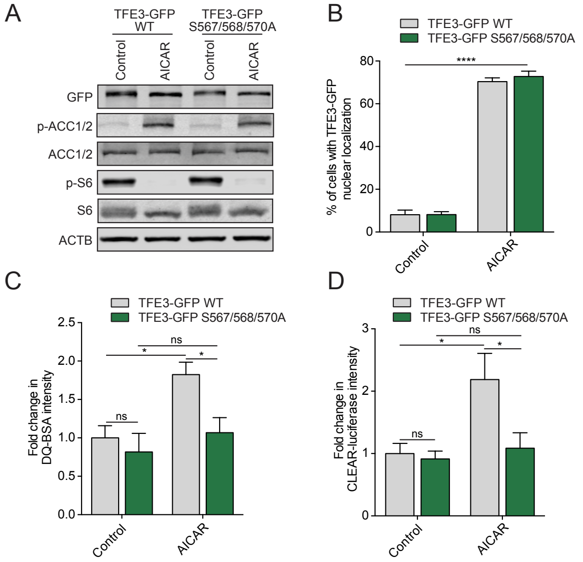


**Figure S3. S567/568/570 phosphorylation is required for TFE3 activity**

(**A**) Immunoblot of protein lysates of MEFs overexpressing TFE3-GFP WT and TFE3-GFP S567/568/570A incubated in complete media (Control) or in presence of AICAR (2 mM) for 2 h. Data are representative of three independent experiments. (**B**) Quantification of the percentage of MEFs overexpressing TFE3-GFP WT and TFE3-GFP S567/568/570A with nuclear TFE3-GFP upon treatments as indicated in (A) (mean ± SEM of three independent experiments, two-way ANOVA, ns=not significant, ****P < 0.0001; n > 200 cells per condition). (**C**) Relative lysosomal activity, as determined by DQ-BSA assay, in MEFs overexpressing TFE3-GFP WT and TFE3-GFP S567/568/570A upon treatment as indicated in (A) (mean ± SEM of three independent experiments, two-way ANOVA, ns=not significant, *P < 0.05; n > 200 cells per condition). (**D**) Relative TFEB/TFE3 transcriptional activity, as determined by CLEAR-luciferase promoter activity normalized against CMV-Renilla in MEFs expressing TFE3-GFP WT and TFE3-GFP S567/568/570A upon treatment as indicated in (A) (mean ± SEM of the luminescence fold change from four independent experiments, two-way ANOVA, ns=not significant, *P < 0.05).

Supplementary Figure 4

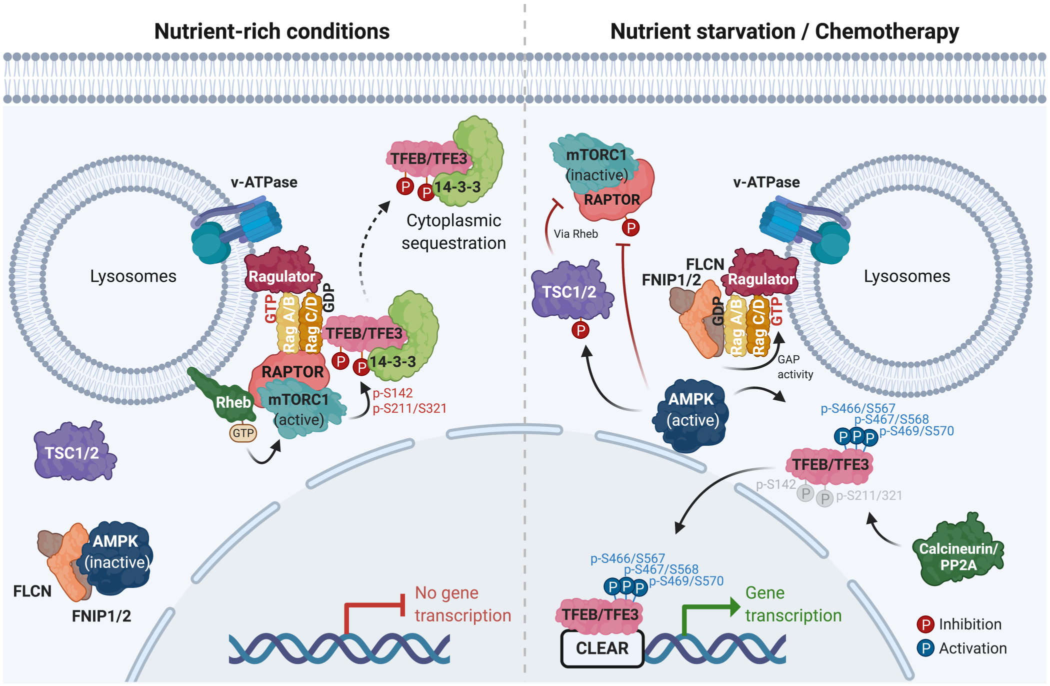


### Figure S4. Model of TFEB regulation by AMPK and mTORC1

Under normal conditions, AMPK is inactive, bound to FLCN/FNIP2, and mTORC1 phosphorylates TFEB and TFE3 on the lysosomes, preventing their nuclear translocation by promoting the binding of 14-3-3. Under starvation or chemotherapy, AMPK is activated and mTORC1 inhibited via TSC2 and Raptor phosphorylation. Calcineurin and PP2a dephosphorylates the mTORC1 sites, promoting TFEB/TFE3 nuclear localization. AMPK phosphorylates TFEB and TFE3 on the serine cluster S466, S467, S469 (S567, S568, S570 for TFE3), which further increases TFEB/TFE3 activity. Created with BioRender.com.
